# Perinatal cannabidiol exposure reshapes astrocyte morphology and tripartite synapse organization in a sex-dependent manner

**DOI:** 10.64898/2026.04.16.719010

**Authors:** Gabriel Henrique Dias de Abreu, Jeremy Wilson, Vitor Ritzmann, Oscar Moosbrugger, Clare T. Johnson, Ken Mackie, Heather Bradshaw, Jui-Yen Huang, Hui-Chen Lu

**Affiliations:** Gill Institute for Neuroscience, Indiana University, Bloomington, IN 47405; Department of Psychological and Brain Sciences, Indiana University, Bloomington, IN 47405; Program in Neuroscience, Indiana University, Bloomington, IN 47405

**Author notes:** Corresponding Author and Address: Hui-Chen Lu, Multidisciplinary Science Building II, 702 N Walnut Grove Ave, Bloomington, IN 47405, USA. Author Contributions: H.C.L., G.H.D.A., J.Y.H., K.M., and H.B. designed the study; G.H.D.A., J.W., C.T.J. performed the experiments; G.H.D.A., J.W., V.R., and O.M. analyzed data. H.C.L. and G.H.D.A. wrote the manuscript with inputs from J.Y.H. and K.M.

**Keywords:** Cannabis, CBD, astrocytes, synapses, development, perinatal exposure

## Abstract

Cannabidiol (CBD) has recently gained significant public acceptance as a safe therapeutic, contributing to increased use during pregnancy. However, little is known about how maternal CBD exposure impacts fetal brain development. Here, we established a preclinical CBD perinatal exposure (CBD-PCE) model to examine the impacts of CBD on astrocyte morphology in the medial prefrontal cortex (mPFC), a brain region critical for working memory and affective behaviors. Astrocytes play critical roles in maintaining ionic/metabolic homeostasis, neurotransmission, and neurovascular coupling in the CNS. They exhibit highly ramified processes with endfeet surrounding synapses, forming tripartite synapses. We quantitatively assessed the impact of CBD-PCE on astrocyte morphology and the composition of tripartite synapses in mPFC using high-resolution three-dimensional (3D) imaging. Our morphometric analyses revealed that CBD-PCE reduced astrocyte density and increased the number of major branches and whole-cell volume in the mPFC of male, but not female, progenies. Using high-magnification 3D analysis, we found that mPFC astrocytes after CBD-PCE exhibited increased neuropil infiltration volume and reduced surface-to-volume ratios in males but not in females. Moreover, the levels of aquaporin-4 (AQP4) and Kir4.1 inwardly rectifying potassium channel, two key components in regulating ionic homeostasis, are elevated on the membranes of male CBD-PCE astrocytes. We also analyzed mPFC tripartite synapses and observed significant increases in thalamocortical tripartite synapse density in both sexes, whereas intracortical excitatory synapses were reduced only in females. Collectively, these findings demonstrate that CBD-PCE induces sex-specific changes in astrocyte morphology and in the composition of tripartite synapses in the mPFC of the progeny’s brains.

## INTRODUCTION

The increasing availability of cannabis-derived products [1], coupled with rising self-medication practices [2], has contributed to increased use of these products by pregnant women [3–5]. The two most studied phytocannabinoids (pCB), Δ^9^-tetrahydrocannabinol (THC) and cannabidiol (CBD), can cross the placenta and accumulate in milk, thereby gaining access to the fetal and neonatal brain [6–9]. While the impact of THC on various aspects of brain development has been better characterized [10–12], little is known about the impacts of CBD on these developmental processes. The consumption of CBD products during pregnancy is often perceived as a “natural” and less-risk approach for the management of nausea, insomnia, anxiety, and chronic pain [13–15] during pregnancy due to the lack of psychomimetic effects [16]. Despite this perception, we and others have employed mouse perinatal cannabinoid exposure (CBD-PCE) models and found that perinatal CBD has lasting effects on behaviors in adult offspring, often in a sex-dependent manner [8, 17, 18]. However, the cellular mechanisms contributing to the impact of perinatal CBD on later behaviors remain to be elucidated.

CBD is a promiscuous molecule and can modulate many signaling pathways, including those of endocannabinoid (eCB) and 5-hydroxytryptamine (5-HT, known as serotonin) [16]. Actions include acting as a negative allosteric modulator of cannabinoid receptor 1 (CB_1_R) [19, 20] and an activator of 5-HT1_A_ receptors [21–23]. The roles of the eCB and 5-HT systems in brain development have been well documented [10, 24, 25] and include glial maturation [26–28]. Astrocytes in the central nervous system (CNS) play critical roles in brain development and function, e.g., synaptogenesis, neurotransmission, neurovascular coupling, and metabolic homeostasis [29]. Astrocytes are mostly born from radial glia cells after neurogenesis [30] and are responsive to environmental changes [31, 32]. In mice, during early postnatal development, astrocytes migrate into the cortex [33] and develop major branches and highly ramified processes. These processes establish largely non-overlapping territorial domains through a process known as tiling [34]. Astrocytic endfeet extensively ensheathe synapses and capillaries, contributing to tripartite synapses and to the structural and functional integrity of the blood-brain barrier [35–38]. Studies have shown that astrocytes are heterogeneous in molecular, morphological, and physiological features, and their morphogenesis is modulated by local neural circuits [39–41]. An important function of astrocytes is the control of extracellular potassium and extracellular space volume. Coordinated water and ion fluxes involving aquaporin-4 (AQP4) and inwardly rectifying potassium channels (Kir4.1) regulate extracellular potassium levels, thereby modulating astrocyte volume. Impairment of this mechanism leads to astrocyte swelling, a phenomenon observed in many pathological conditions [42–46].

To examine the cellular effects of CBD, we first established a translationally relevant CBD-PCE model that mimics CBD exposure across all three trimesters of human pregnancy. Given the critical role of astrocytes in neural circuit function, we quantitatively assessed the impact of PCE on astrocyte morphogenesis and tripartite synapses. Using high-resolution three-dimensional (3D) imaging and detailed morphometric analyses, we found that CBD-PCE induces sex-dependent alterations in astrocyte density, astrocyte morphogenesis, AQP4/Kir4.1 abundance, and tripartite synapse density. Together, these findings provide the first evidence that CBD-PCE affects astrocyte morphogenesis and tripartite synapse organization.

## MATERIALS AND METHODS

### Animals

Mice were housed 4 or 5 per cage under a 12h light/dark cycle, with food and water provided *ad libitum*. All animal breeding and experimental procedures were approved and performed in accordance with the guidelines of the U.S. Department of Health and Human Services and the Indiana University Bloomington Institutional Animal Care and Use Committee. We used the outbred CD1 mouse strain because its genome heterogeneity is similar to that of wild-caught mice [47] and it harbors fewer mutations than those of commonly used inbred strains.

Additionally, CD1 dams exhibit robust breeding and mothering behaviors. Aldh1l1-GFP mice in a C57BL/6J background [48, 49] were kindly provided by Benjamin Deneen (Baylor College of Medicine). These reporter mice express cytoplasmic GFP driven by the Aldh1l1 pan astrocyte promoter [48] and thus all astrocytes in the brain are labeled. Aldh1l1-GFP mice were backcrossed into CD1 by crossing Aldh1l1-GFP heterozygous mice with CD1 mice for >5 generations. Experimental groups were composed of mice from 3 or 4 different dams to minimize litter effects.

### Drugs and injections

Δ^9^-tetrahydrocannabinol (THC), and cannabidiol (CBD) were provided by the Drug Supply Program, National Institute on Drug Abuse (NIDA). The combination of THC+CBD was prepared with the dilution of CBD powder with an equimolar amount of a 318 mM ethanol stock solution of THC. Stock solutions were prepared in 100% ethanol and serially diluted to obtain a 25 mM stock, from which injection solutions were freshly prepared. Daily injection solutions were prepared in a vehicle consisting of Cremophor® EL (Millipore-Sigma, cat# 238470), 100% ethanol, and sterile saline in a ratio of 1:1:18, respectively, and administered at a volume of 10 mL/kg. Timed pregnancies were established with trios (1 male with 2 females), and the morning when a vaginal plug was found was denoted as gestational day 0 (GD0). The male was removed from cage at this day. From GD5 to GD20-21 dams were injected subcutaneously (s.c.) daily between 2 and 5 pm with vehicle or 3 mg/kg of THC, CBD or THC+CBD. From postnatal day 0 (P0) to P10 (P10) pups were injected with a BD SafetyGlide 29G syringe with 1 mg/kg s.c.

### Development of the time and dosage-informed PCE model

The selected dosing strategy was based on prior reports demonstrating that 3 mg/kg pCB administration in mice yields plasma concentrations within the human psychoactive range [50], as well as on our previous findings of comparable pCB levels reaching embryonic brains following maternal administration [8] and pCB accumulation in milk [9]. To assess pCB transferred from maternal milk to pup P10 cortices, nursing dams received daily s.c. injections of 3 mg/kg of THC+CBD from P0 to P10. Pups were allowed to feed freely. At P10, cortices were collected 2 hours after the last dam injection and flash frozen in liquid nitrogen until mass spectrometry (MS) measurements. Next, two independent cohorts were treated daily with either THC or CBD (Figure 1A). First, pregnant dams received s.c. injections of 3 mg/kg of THC or CBD from GD5.5 to 18.5. Secondly, to quantify pCB accumulated in P10 cortices after direct pup injection, samples received via s.c. injections of 1 mg/kg or 3 mg/kg, of THC or CBD, from P7-P10. 2 hrs after the last injection, cortices were quickly dissected and flash frozen in liquid nitrogen until MS measurements. The data points for pCB levels in E18.5 brains are the same as previously published by our group [8] and now used for comparison with P10 pCB analysis.

**Figure 1.**
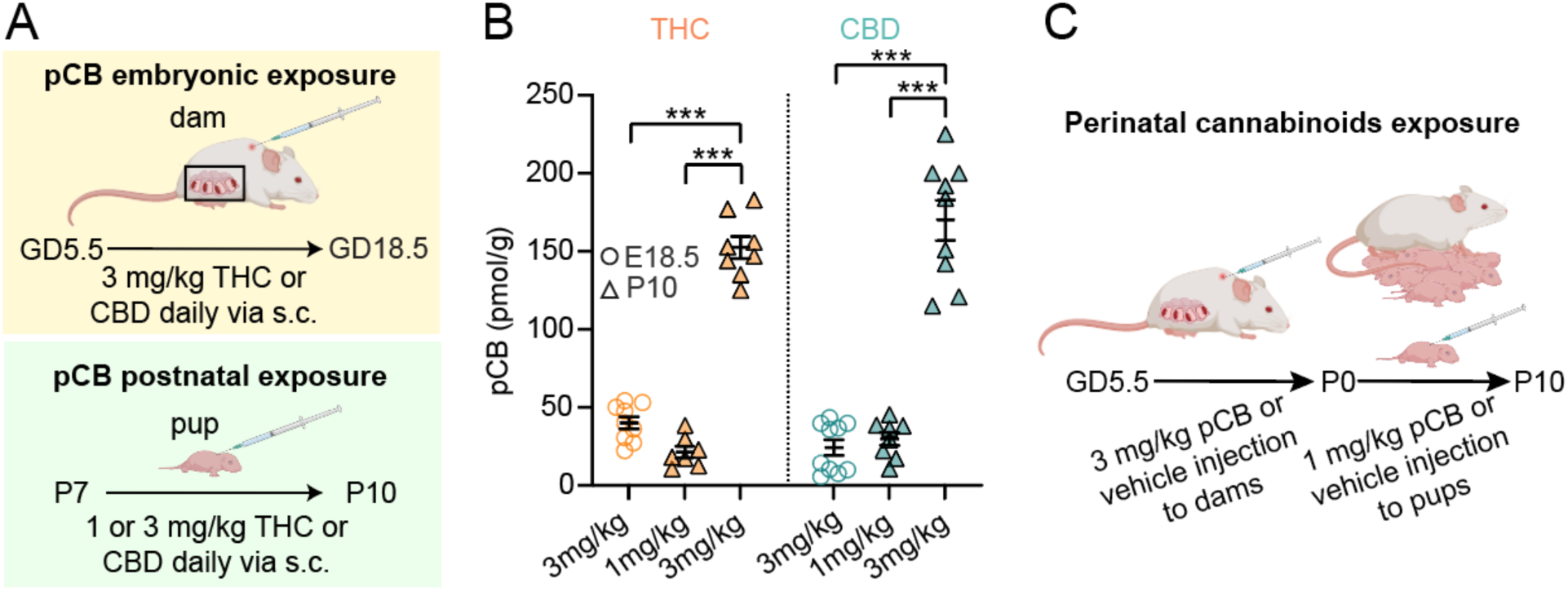
Brain THC and CBD levels 2 hrs after the final pCB injection. (A) Timelines of embryonic and postnatal pCB exposure paradigms via different routes of administration. For embryonic exposure (top), dams were treated via subcutaneous (s.c.) injections daily from GD5.5-18.5. For postnatal exposure (bottom), pups were injected s.c. daily during P7-10. (B) Summary for brain pCB levels. (C) Schematic illustrates the pCB-PCE treatment regimen used in this study. Two-way ANOVA, dose × cannabinoid, Tukey’s post hoc. Each data point represents an individual animal, and error bars represent mean ± SEM. E18 embryo numbers: n=9 for THC; n=10 for CBD. E18.5 datapoints previously published [8]. P10 pup numbers: n=7 for 1 mg/kg THC; n=8 for 1 mg/kg CBD, n=8 for 3 mg/kg THC; n=9 for 3 mg/kg CBD. *p ≤ 0.05, **p ≤ 0.01, ***p ≤ 0.001.

### Lipid extraction/HPLC/MS/MS phytocannabinoid analysis

E18.5 embryo brain and P10 cortices were harvested 2 hours after the last cannabinoid injection, and samples were flash frozen in liquid nitrogen and stored in -80 °C. On the day of lipid extraction, samples were mixed with 2 mL of 100% methanol, and 5 µL of 1 µM deuterium-labeled AEA (AEA_d8_ – Cayman Chemical, cat# 390050) were added as described [9, 51, 52]. Samples were incubated on ice in the dark for 30 min before being sonicated and centrifuged at 19,000 x *g*, 20 °C, for 20 min. The resulting lipid-containing supernatants were decanted into 7 mL of water to create a 25% organic solution. This solution was then partially purified on C18 solid-phase extraction columns as previously described [51], and lipids were eluted with 1.5 mL of 3 different methanolic concentrations (65, 75, and 100%). Elutions were stored at -80 °C until high-performance liquid chromatography-tandem mass spectrometry (HPLC/MS/MS) analysis using an Applied Biosystems API 3000 triple quadrupole mass spectrometer with electrospray ionization (Foster City, CA, USA) in multiple reaction monitoring (MRM) mode.

Analytes were identified by retention time and mass-to-charge ratio, compared with known standards. Analyst software (Applied Biosystems) was used to determine analyte concentrations, as previously described [9, 51, 52]. Concentrations were converted to moles per gram for each sample and corrected for extraction efficiency based on the percent of the internal standard, AEA_d8_, in each individual sample. The samples in which the concentration of an analyte deviated by more than 2 standard deviations from the group mean were excluded from analysis.

### Genotyping

Ear tissue from individual mice was lysed in tissue digestion buffer (50 mM KCl, 10 mM Tris-HCl, 0.1% Triton X-100, 0.1 mg/mL proteinase K, pH 9.0), vortexed gently, and then agitated overnight at 600 rpm at 55 °C. Samples were then heated to 95°C for 20 minutes to denature proteinase K (Thermo Scientific, Rockford, IL, USA). Supernatants of the samples after centrifugation at 16,100 x *g* for 15 minutes were used as DNA templates for polymerase chain reactions (PCRs, EconoTaq Plus Green 2x Master Mix, Lucigen, Middleton, WI, USA). The primer sequences for Aldh1l1-GFP were: Forward-5’-CCTCTGGCTGCTCCTTCAACAG-3’, reverse-5’-GGTCGGGGTAGCGGCTGAA-3’, and common-5’-CCTTAGCTGGCAGTAAACCTCCTG-3’.

### Immunostaining

Mice were deeply anesthetized and perfused with 4% paraformaldehyde (PFA) in PBS using a peristaltic pump at ∼3 mL/min for 10 min. Brains were harvested and postfixed with 4% PFA/PBS overnight at 4 °C. Brain sections were obtained using a vibratome (VT-1000 Vibrating Microtome, Leica Microsystems) and stored in antigen preservation solution (NaCl 137 mM, KCl 2.7 mM, Na_2_HPO_4_ 10 mM, KH_2_PO_4_ 1.8 mM, 50% (v/v) ethylene glycol and 1% (w/v) of polyvinylpyrrolidone) at -20 °C until staining. For cell counting and Aldh1l1-GFP processes reconstruction, we used 50 μm coronal brain sections; for other analyses, we used 100 µm sections. First, coronal sections were quickly rinsed in 1x PBS to remove the storage solution, permeabilized with 2% Triton-X100 in PBS for 30 min at room temperature and incubated with blocking solution (3% normal goat serum, Jackson ImmunoResearch) in 2% Triton-X100 in PBS for 1 hour at room temperature. Primary and secondary antibodies were prepared in blocking solution and incubated for 48 hours at 4 °C. Detection of primary antibodies was performed with appropriate secondary antibodies conjugated to Alexa Fluor dyes. Cell nuclei were labeled with DAPI (100 ng/mL, ThermoFisher) added to the secondary antibody solution. Sections were mounted in ProLong Gold Antifade mounting medium (ThermoFisher, cat# P36934) and stored in the dark for 48 hours before imaging. Supplementary Table lists the antibodies used.

### Cellpose2.0-assisted cell counting analysis

Confocal microscopy images were acquired from the mouse prefrontal cortex (bregma 1.98mm to 1.54mm) [53] using a Nikon Ti-2 A1 confocal microscope equipped with a 10x objective (NA 0.45) at 2048 × 2048 pixel resolution. Four-channel sequential z-stacks (DAPI, Aldh1l1-GFP, NeuN, and S100β) were acquired at 1 µm z-step intervals, resulting in a total thickness of 5 µm. For post-acquisition processing, all images were z-average projected, and all channels were cropped to the prelimbic region of prefrontal cortex based on the NeuN staining pattern using Fiji-ImageJ. A rectangular region-of-interest (ROI) with fixed width (317 µm) and variable length was defined to span the entire cortical depth, extending from the bottom of layer 6 to the top of layer 1. Single channel images corresponding to individual cellular markers were exported from the ROI-defined regions for Cellpose-assisted quantification. Cell counting was performed using a human-in-the-loop approach with Cellpose2.0 [54], with an experimenter blinded to treatment conditions. Initial segmentation was performed using the pretrained “cyto3” model for all cell types. These outputs were manually corrected to generate ground-truth datasets, which were subsequently used to train custom models for each cell type. Custom-trained models were fine-tuned to enhance the detection of specific markers (Aldh1l1-GFP, NeuN, S100β). Cell diameter parameters were automatically estimated for each marker: GFP (11.3 µm), S100β (10.57 µm), and NeuN (14.83 µm). Segmentation performance was further optimized by adjusting the model-specific parameters: cellprob_threshold and flow_threshold were set to -0.9, 0.7 for GFP, -1.2 and 1.0 for S100β, and -1.0 and 0.6 for NeuN, respectively. Custom models were trained for 300 epochs with a learning rate of 0.3 using approximately 150-200 ROIs per cell type (derived from ∼2-3 images per marker). The resulting models enabled reliable identification of neurons and astrocyte populations. Cellpose segmentation outputs were visually inspected, and cellular masks were manually corrected where necessary to ensure accuracy. For spatial quantification across cortical depth, each ROI was subdivided into 10 equal bins along the radial axis (from layer 1-6). The number of segmented cells within each bin was quantified using custom Python scripts, enabling the generation of cell distribution profiles across cortical layers.

### Astrocytes imaging acquisition for process tracing and analysis

To quantify astrocyte branching complexity, astrocytes were imaged from 50 µm-thick coronal brain sections of Aldh1l1-GFP mice encompassing the prefrontal cortex (bregma 1.98mm to 1.54mm) [53], specifically within layer 5 of prelimbic region. Imaging was performed using a Nikon Ti-2 A1 confocal microscope equipped with a 60x oil-immersion objective (NA 1.4) at 1024 × 1024 pixels resolution. Z-stack images were acquired at 0.5 µm intervals spanning the full thickness of each section. Image stacks were imported to Imaris software (v9.8.2; Oxford Instruments) for three-dimensional (3D) reconstruction of astrocyte morphology. Only astrocytes with somata fully contained within the z-stack and exhibiting intact, clearly discernible processes were selected for analysis. To enable the automatic tracing of major astrocytic branches using Filament module (Autopath mode), a two-step processing workflow was applied. First, the brightest GFP signal corresponding to the soma and primary processes was segmented using the Surface module with auto-thresholding applied on a per-image basis. These surface-rendered objects defined individual astrocyte regions. Second, a new channel was generated by retaining only the fluorescent signal within the segmentation surface objects, thereby restricting the signal to a single astrocyte. Astrocyte processes were reconstructed using the Autopath mode in Filaments module. Soma seeds were defined by placing spherical starting point with a diameter of 10 µm, from which filament tracing was initiated. For each reconstructed astrocyte, total filament length, number of branching points, and number of Sholl intersections (radius set at 5 µm) were quantified as measured for morphology complexity. All image processing, segmentation, and tracing procedures were performed by experimenters blinded to experimental conditions.

### Sparse labeling of astrocytes

Astrocytes were sparsely labeled in CD1 mice PCE offspring using AAV5.GfaABC1D.Lck-GFP.SV40 (Addgene #105598-AAV5) via free-hand intracerebroventricular (ICV) injections at P0, as previously described [55]. Briefly, neonatal pups were anesthetized by hypothermia and received bilateral injections of 1 µL AAV (1 × 10¹¹ vg/mL in sterile 1x PBS + 0.05% fast green dye) into the lateral ventricles using a Hamilton syringe (32G, cat. #7803-04, RN 6PK PT4, 0.375”). Following injections, pups were allowed to recover on a heated pad for 10 minutes before being returned to the dam. Brains were collected at P21 and processed for subsequent immunolabeling analyses.

### Astrocytes whole-volume and neuropil infiltration volume imaging and analysis

Whole-astrocyte images were acquired with a Nikon Ti-2 A1 confocal microscope with a 60x oil objective (NA1.4) at 1024 × 1024 pixel resolution, and pixel size: 0.1437 µm x 0.1437 µm x 0.5 µm (x, y, z), and 2x digital zoom to sample one astrocyte per image. DAPI and GFP channels were imaged sequentially at 0.5 µm z-step intervals from top to bottom. Astrocytes were sampled from layers 5 or 6 of the prelimbic region in the prefrontal cortex (bregma +1.98mm to +1.54mm) [53] across 2 or 3 brain sections from both hemispheres. Z-stacks were analyzed using Imaris software (v10.0.0; Oxford Instruments) following a workflow adapted from previously published methodologies [56, 57].

First, the Surface module was used to render objects of nuclei and vasculature-like artifacts. We created a new channel where these structures were masked out from the GFP channel to exclude their occupied volume. We applied the Surfaces module to the previously cleaned GFP channel to generate full astrocyte volumes and surface areas. To assess neuropil infiltration volume (NIV) from astrocyte membranes, in Surface module we positioned four identical ROI volumes of 5.75 µm x 5.75 µm x 2 µm (x, y, z) distributed across the z plane in areas deprived of astrocyte primary branches but in proximity to DAPI signal, and rendered the volume occupied by GFP signal in the ROI volume. Data were normalized to % of ROI volume occupied. A graphic workflow for representative images of NIV assignment workflow is provided in Supplementary Fig. 4A-B.

### AQP4 and Kir4.1 density on astrocyte membrane colocalization imaging and analysis

Lck-GFP-labeled astrocytes were co-stained with AQP4 and Kir4.1 antibodies for analysis of astrocytic-membrane water and inward rectifying K^+^ channels, respectively. To obtain an initial qualitative assessment of GFP-tagged astrocytic membrane organization under diffraction-limited confocal imaging conditions, whole astrocytes spanning 35-40 µm in the z-axis were imaged using four sequential channels acquired in two alternating sweeps (DAPI-AQP4/GFP-Kir4.1). Imaging was performed on a Nikon Ti-2 A1 confocal microscope with a 60x oil objective (NA1.4) at 1024 × 1024 pixels resolution using 3x digital zoom and 2x frame averaging, with a pixel size of 0.096 µm x 0.096 µm x 0.1 µm (x, y, z), providing lateral and axial sampling that oversampled the diffraction-limited resolution to support better 3D visualization and postprocessing deconvolution. Acquisition of a full astrocyte z-stack required approximately 1h10min (Supplementary Fig. 3A).

For the AQP4 and Kir4.1 dataset, post-acquisition raw files were deconvolved in NIS Elements Analysis AR using the 3D Lucy-Richardson algorithm. Because both markers formed fine, membrane-enriched microdomains along the astrocytic membrane tagged with GFP, accurate detection required improved axial resolution and reduced out-of-focus blur to prevent signal spreading into adjacent structures. To ensure reliable quantification of these small, membrane-confined puncta, the raw images were, therefore, deconvolved prior to segmentation. Processing parameters were identical across all samples and matched to the imaging configuration: objective NA1.4, refractive index: 1.515, pixel size: 0.096 µm, and z-step 0.1 µm. A theoretical point spread function (PSF) was generated automatically using the excitation wavelengths (561 and 640 nm) and corresponding emission detection ranges. The algorithm executed 60 iterations, with chromatic shift correction enabled and default background suppression. All deconvolved stacks were visually inspected to confirm the absence of processing artifacts. The deconvolved channels were subsequently used for quantitative spot analysis to evaluate overall puncta density and membrane-associated expression of AQP4 and Kir4.1.

During data acquisition and analysis, our volume inspection along the z-axis showed that the Lck-GFP-labeled astrocytic membrane exhibits complex membrane-associated patterns within the imaged volume, forming intricate structures that tile neighboring cell nuclei and parenchymal elements, including at least one major blood vessel delineated by AQP4+ astrocytic endfeet (Supplementary Fig. 3A-B). Along these GFP-positive membrane domains, discrete, spot-like AQP4- and Kir4.1-immunoreactive signals were consistently observed, consistent with clustered membrane-associated channel expression [42, 58, 59] (Supplementary Fig. 3B-D). This labeling pattern remained consistent across z-stacks encompassing the astrocyte soma when acquired using identical imaging parameters. Consequently, we adapted our acquisition parameter to a total z-depth of 5 µm centered on the astrocyte soma (Supplementary Fig. 3C), allowing optimization of imaging time while preserving representative membrane-associated signal detection.

AQP4 signal associated with astrocyte endfeet, and vasculature was excluded using a multi-step masking workflow implemented in Imaris (v10.2.0; Oxford Instruments). A graphic of the workflow is presented in Supplementary Fig. 4E-F. First, DAPI and vasculature-like signals were rendered using a machine learning–based algorithm with the Surface module and masked out from the AQP4 and Kir4.1 channels. The previously generated AQP4 channel was then used to generate an integral surface object defining the analyzable neuropil volume. In deconvolved datasets, we analyzed AQP4-immunoreactive puncta forming discrete, spot-like signals along the astrocytic membrane. The anti-AQP4 antibody used was generated against a recombinant immunogen corresponding to AA 249-323 of mouse AQP4 isoform 2 (UniProt ID: P55088-1), corresponding to the AQP4-M1 isoform. Given the diffraction-limited resolution and the labeling of non-orthogonal array-forming by this isoform [59–61], we employed an object-based analysis approach using Imaris to estimate the overall density of AQP4 signal rather than individual cluster size. The same analysis pipeline was applied to Kir4.1 expression assessment.

AQP4 and Kir4.1 puncta were quantified using the Spots module. First, we normalized detection of “Quality %” parameter threshold across the entire dataset for each marker. Thresholds were selected to maximize puncta detection without introducing artifacts and were applied uniformly, with spots diameters set to 0.3 µm for both markers [60]. We estimated spot size by navigating through the z-stack using slice view mode in Imaris, measuring the lateral distance between opposing edges of individual puncta at the plane of maximal intensity, and determining the number of z-planes over which the same discrete signal persisted to confirm 3D continuity. The overall number of spots for AQP4 and Kir4.1 were extracted from spots detected in the neuropil object using Imaris MATLAB R2025a XTension “Split spots into surface” and normalized to the neuropil volume. In parallel, we also generated a Surface-based object corresponding to astrocyte membrane-bound GFP expression.

Lastly, previously detected Spots for AQP4 and Kir4.1 in the volume were applied to “Find spots close to surface” with the astrocyte GFP surface defined as reference and distance threshold set to zero. This workflow allowed us to estimate the number of spots corresponding to each marker at the astrocyte surface. Data were extracted and expressed as the overall densities of AQP4 and Kir4.1 puncta. Spot density values were subsequently normalized to the astrocyte surface area to account for differences in cell size captured.

### Synaptic markers density within astrocyte neuropil imaging and analysis

Sparsely Lck-GFP-labeled astrocytes were co-stained with synaptic markers to identify putative excitatory and inhibitory synapses. For excitatory synapses, the intracortical-vesicular glutamate transporter 1 (VGLUT1) and thalamocortical-vesicular glutamate transporter 2 (VGLUT2) [62, 63] presynaptic markers were used in combination with the postsynaptic marker, postsynaptic density 95 (PSD95). For inhibitory synapses, vesicular GABA transporter (VGAT) [64, 65] was used as the presynaptic and gephyrin as the postsynaptic marker. Three sets of staining (DAPI+GFP+VGLUT1+PSD95; DAPI+GFP+VGLUT2+PSD95; DAPI+GFP+VGAT+Gephyrin) were imaged individually at P21 with the same brain coordinates and high-resolution imaging acquisition parameters as detailed above.

The analysis workflow was adapted from previous published methodologies [66–71]. The volume occupied by nuclei and vasculature varied across images, substantially affecting the density of synaptic markers quantification. To better estimate the neuropil volume in which the density of synaptic markers were quantified, we first rendered surface objects of nuclei and vascular-like objects employing a machine-learning surfaces algorithm and then masked these objects from pre- and postsynaptic marker channels. The post-masked PSD95 or gephyrin channel was used to generate an integral surface object containing the total analyzable volume (neuropil object).

Presynaptic (VGLUT1, VGLUT2, and VGAT) and postsynaptic (PSD95 and gephyrin) punctuate were detected using Imaris Spots module with 0.5 and 0.3 µm diameter threshold, respectively. We estimated spot size by navigating through the z-stack using slice view and measured the distance between points spanning a punctum and determining the number of z-planes over which the same spot signal persisted. These measures were consistent with previously reported synaptic puncta diameters ranging from 0.25 to 0.8 µm [72, 73].

To determine optimal Spots detecting thresholds, we manually normalized the detection “Quality %” parameter across the entire dataset for each synaptic marker, selecting a threshold that maximized puncta detection while introducing no artifacts. This uniform threshold was then applied to all images. We employed an object-based juxtaposition workflow using Imaris MATLAB R2025a XTension “Split spots into surface” to quantify Spots density corresponding to each synaptic marker inside the neuropil volume. Next, we applied XTensions “Colocalization” to previously detected spots to determine presynaptic directly opposed to postsynaptic spots, representing anatomically defined putative synapses within 0.5 µm distance threshold.

To segment the tripartite synaptic environment, with the Surface module we applied 4 ROIs of 5.75 µm x 5.75 µm x 5 µm (x, y, z) encompassing the 50 z planes, positioned within the NIV devoid of primary astrocytic branches, rendering GFP volume corresponding to the astrocytic membrane occupying each fixed ROI in the NIV. Finally, to find previously colocalized Spots corresponding to pre and post synaptic markers within astrocyte NIV, we applied “Find spots to surface” function from Imaris XTensions module, using a 0.5 µm distance threshold from the previously annotated juxtaposed spots to the rendered GFP surface in each ROI. Data were extracted and normalized as the density of juxtaposed spots relative to the density of GFP volume within the ROI volume. A graphic of the workflow is shown in Supplementary Fig. 4C-D. The 3D analysis workflow was adapted from Stogsdill, et al; and Irala, et al. [56, 74].

### Statistical analysis

Mass spectrometry (MS) data used for PCE model development (Figure 1 and Supplementary Fig. 1) were analyzed using GraphPad Prism 10. Individual data points represent biological replicates (n = 1 animal), and results are presented as mean ± SEM. Differences across cannabinoid type (THC vs. CBD) and dose were evaluated using a two-way analysis of variance (ANOVA) with interaction terms, followed by Tukey’s post hoc multiple comparisons test. Statistical significance was defined as p < 0.05, with thresholds indicated as *p < 0.05, **p < 0.01, and ***p < 0.001. Data in Supplementary Fig. 1 are presented descriptively to illustrate central tendency and variability and were not subjected to inferential statistical testing.

Histological data in this study exhibited a clustered structure (e.g., cells nested within regions of interest, and regions within animals) [75–77]. To appropriately account for this hierarchical organization and ensure accurate estimation of statistical significance, multilevel-statistical analyses were performed as described previously [78, 79] In brief, datasets were primarily analyzed using linear mixed-effects (LME) models to account for within-subject and between-subject variability across the reported measurements. When datasets did not exhibit hierarchical or nested organization, standard linear regression (LM) were used instead. Model refinement was performed by through stepwise exclusion of terms that did not improve model performance, as evaluated using Aikake Information Criterion and Bayesian Information Criterion metrics. To obtain robust and unbiased estimates of variance components for both LME and LM outputs, we employed a bootstrapping approach to resample the dataset (n=10,000) to simulate the underlying data distribution. The resulting bootstrapped estimates were then used to perform post-hoc pairwise comparisons, which served as the basis for the final reported statistics. Description of individual data points, whether representing images, cells, or NIV ROIs, is provided in each figure legend. Data and analysis materials supporting the findings of this study are available from the corresponding author upon reasonable request.

## RESULTS

### Establish a clinically relevant perinatal cannabis exposure (PCE) treatment regimen

We first aimed to establish a mouse model of pCB exposure that mimics human cannabis use during pregnancy, in which maternal consumption leads to indirect exposure of the developing offspring. Because altricial rodents are born at a relatively immature stage of brain development, early postnatal mouse brain development corresponds to the third trimester of human fetal brain development [80, 81]. Because the human cortex at birth is developmentally similar to the mouse cortex at postnatal day 10 (P10), it is necessary to extend pCB exposure into the early postnatal period. Our previous work shows that maternal milk contained THC and CBD after the nursing dam received THC+CBD subcutaneous (s.c.) injections [9]. However, we found that much less CBD compared to THC delivered via lactation accumulates in pup brains, suggesting limited absorption from the pup’s gut (Supplementary Fig. 1).

To establish a pCB dosing regimen that offers comparable exposure across embryonic and early postnatal stages, we treated two independent cohorts daily with either THC or CBD (Figure 1A). In the first cohort, pregnant dams received subcutaneous (s.c.) injections of 3 mg/kg of THC or CBD from GD5.5 to 18.5. This dosing strategy was based on previous reports showing that 3 mg/kg pCB administration to mice yields plasma concentrations within the human psychoactive range [50], as well as on our previous findings demonstrating comparable drug levels in embryonic brains [8]. In the second cohort, pups were administered either 1 mg/kg or 3 mg/kg of THC or CBD from P7-P10. The lower dose was chosen because of low levels of pup pCB-metabolizing P450 enzymes [82]. Brain tissue from E18.5 embryo and P10 pups were collected two hours after the last injection and processed for HPLC/MS analysis to measure pCB brain content. As expected, maternal exposure to THC or CBD was sufficient to induce pCB accumulation in embryonic brains. In the postnatal injection cohort, both THC and CBD accumulate in pup brains in a dose-dependent manner (Figure 1B). Notably, pups administered with 1 mg/kg of THC or CBD exhibited brain levels comparable to those observed in the embryonic brain following maternal exposure to 3 mg/kg.

These data informed the development of a preclinical pCB-PCE model incorporating postnatal dosing to achieve brain concentrations comparable to those observed *in utero* (Figure 1C). In the following experiments, THC- and CBD-PCE pups were generated by daily subcutaneous (s.c.) administration of 3 mg/kg THC, CBD, or THC+CBD to pregnant dams from G5 to birth, followed by daily s.c. injections of 1 mg/kg THC or CBD to pups from P1-P10. The brains of these pups were harvested for examinations at P21, the weaning age, after 11 days free of pCB exposure.

### PCE does not change the mPFC cytoarchitecture but reduces neuron and glia densities

PCE could potentially affect several critical aspects of early brain development, including neuron and glia proliferation and the formation of cortical layers [12, 30, 83, 84]. To explore whether THC-, CBD-, or THC+CBD-PCE alters these early developmental processes, we examined cytoarchitecture and cell density in the mPFC-prelimbic (mPFC-PrL) regions in P21 PCE- and vehicle-control progeny. Aldh1l1-GFP heterozygous mice were selected for this examination because most astrocytes in this transgenic mouse line are labeled with cytoplasmic GFP, providing a great tool to evaluate the impact of PCE on astrocyte distribution and numbers [48] (Figure 2A). The mPFC-PrL region was selected for its known vulnerability to developmental cannabinoid exposure [85–89] and its central role in cognitive and emotional processes, including working memory and anxiety [90, 91] (Figure 2B). Immunostaining with antibodies against NeuN (a neuronal marker) [92], GFP (amplified the astrocytic GFP signals), and S100β (a marker for a subset of astrocytes and oligodendrocyte precursor cells [93, 94]), was conducted with Aldh1l1-GFP brain sections acquired after PCE and vehicle treatment (Figure 2C). The densities and distributions of these neurons and glia across differential cortical layers in the mPFC-PrL were quantified (Figure 2D).

**Figure 2.**
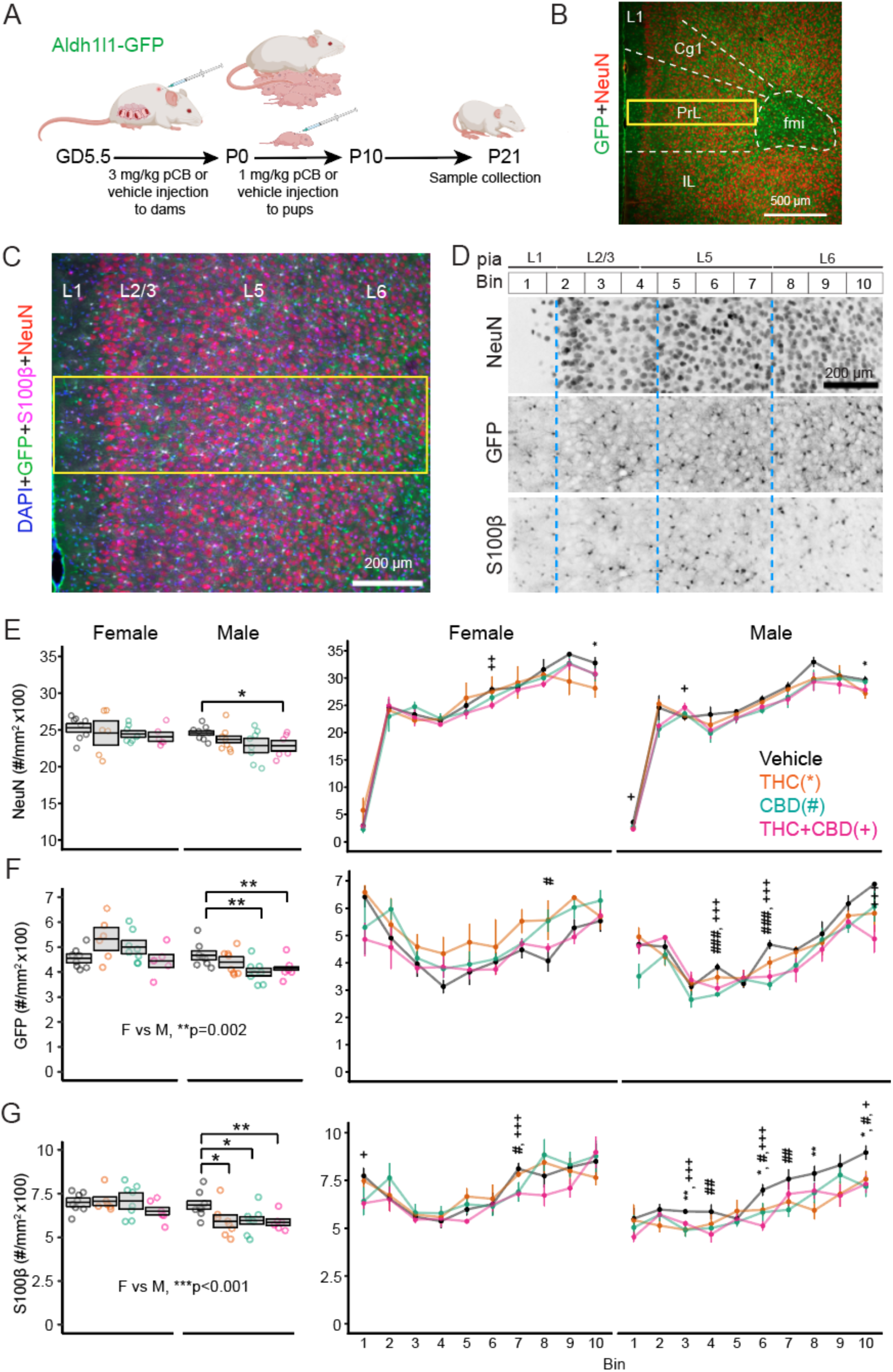
Cellpose2.0-assisted analysis for neuron and glia densities and distributions in mPFC-PrL. (A) Schematic shows the pCB-PCE paradigm with Aldh1l1-GFP mice. (B) The atlas diagram illustrates the brain section containing mPFC-PrL selected for immunostaining experiments (left). A representative image of GFP+NeuN double staining taken from the mPFC region corresponding to the region marked with the red square (right). (C) Representative NeuN, GFP, S100β, and DAPI staining image from the mPFC-PrL region acquired for Cellpose 2.0 analysis. The yellow rectangular box represents the ROI selected for cell counting. (D) Individual channel images from the ROI spanning 10 equal-sized bins from L1 to L6. (E-G) Summary graphs for total density of NeuN, GFP, and S100β, respectively as well as summary graphs of cell densities across bins spanning L1-L6. Numbers of mice and images analyzed per group: Vehicle (4F, 8 images; 4M, 8 images); THC (3F, 6 images; 4M, 8 images); CBD (4F, 8 images; 4M, 8 images); THC+CBD (3F, 6 images; 3M, 6 images). Statistical significance was assessed using multilevel modeling. Data are presented as mean ± SEM derived from bootstrapping output; *p < 0.05; **p < 0.01; ***p < 0.001. Significance symbols for bin analysis, *vehicle vs THC; ^#^vehicle vs CBD; ^+^vehicle vs THC+CBD. Abbreviations: L1, L2/3, L5, L6 for cortical layer1, 2/3, 5, 6, respectively; Cg1, cingulate cortex; fmi, forceps minor of the corpus callosum; PrL, prelimbic cortex; IL, infralimbic cortex.

Our imaging data show that, grossly, cortical layering is normal for both neuronal and glial distributions in the PCE P21 mPFC (Figure 2E-G), except for subtle yet significant distributional differences by PCE. Regarding total cell densities, we found that THC+CBD-PCE significantly reduced total neuron density in the mPFC-PrL of male but not female offspring (Figure 2E). Both CBD- and THC+CBD-PCE reduced total astrocyte densities in male but not female offspring (Figure 2F). The densities of S100β^+^ cells were significantly reduced only in male progenies of THC-, CBD-, and THC+CBD-PCE (Figure 2G). These cell distribution and density analyses reveal that PCE results in significant reductions in both neuron and glia densities in the mPFC of male progeny.

### CBD-PCE increases astrocyte branching complexity in male but not female mPFC-PrL

Astrocytes are responsive to environmental changes and develop their ramified processes into largely non-overlapping territorial domains known as tiling [31, 32, 34, 84]. The observation of reduced astrocyte densities in mPFC-PrL by CBD- and THC+CBD-PCE motivated us to quantitatively evaluate PCE impacts on astrocyte morphologies. The cytosolic Aldh1l1-GFP expression labels major branches of astrocytes [48, 49]. Thus, we used Aldh1l1-GFP mice to examine whether PCE alters the major branch patterns of astrocytes in the mPFC-PrL layer 5 (L5) region, where we observed the greatest changes in astrocyte density (Figure 3).

**Figure 3.**
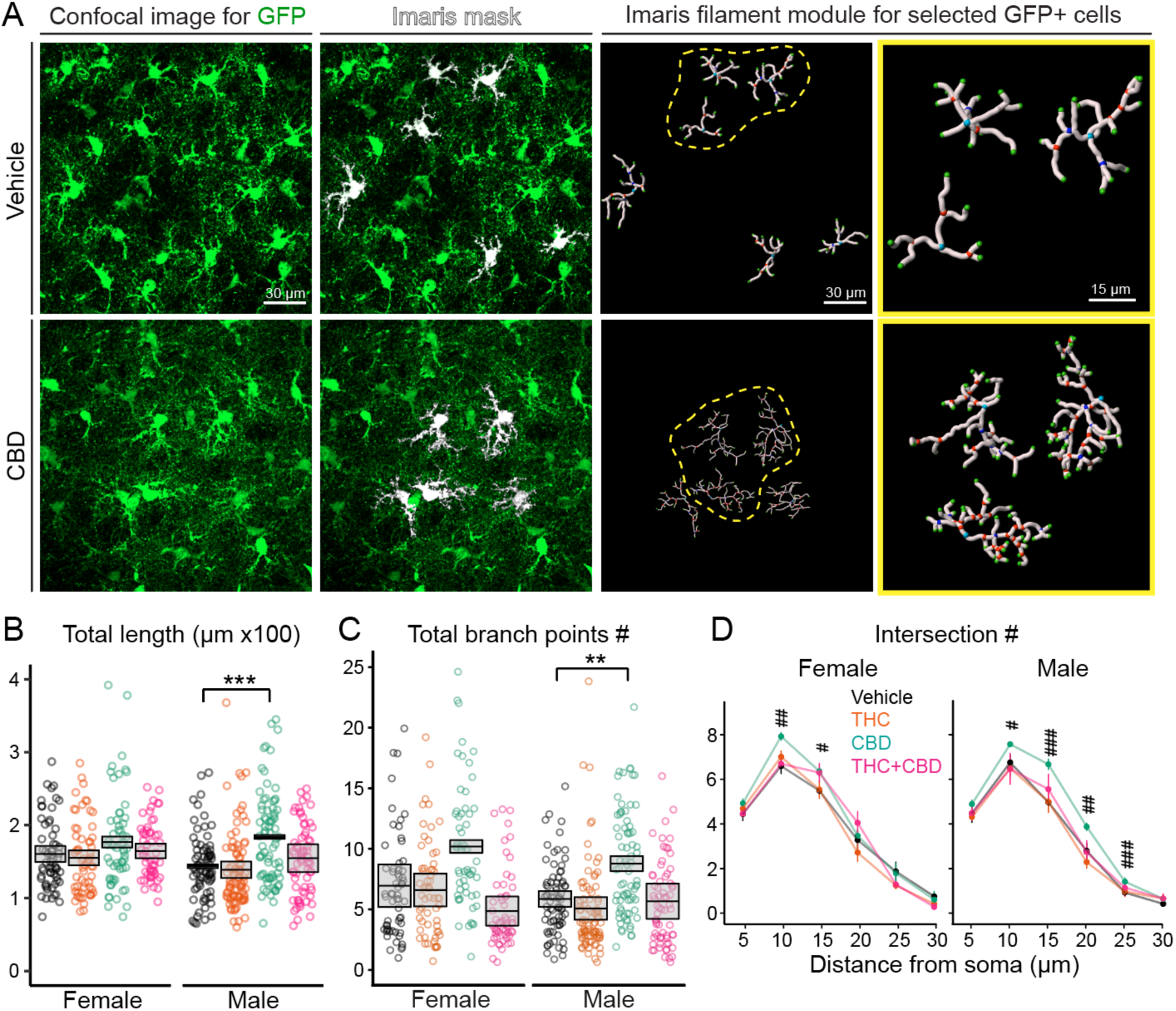
Primary branch pattern analysis with L5 mPFC astrocytes labeled by Aldh1l1-GFP to examine pCB-PCE impacts. (A) Representative confocal images of Aldh1l1-GFP astrocytes from vehicle and CBD-PCE male mice at P21. Imaris masks (white) highlight selected GFP^+^ astrocytes for reconstruction using the Imaris filaments module to label major processes and branching points. Sky blue, soma; orange, branching points; green, ending points. Yellow outlines the enlarged view of selected astrocytes (dashed yellow lines). (B-C) Summary graphs for total length (B) and branching points (C). (D) Sholl analysis for branch intersections at different distances away from the soma; ^#^vehicle vs CBD. Numbers of mice and cells analyzed per group: Vehicle (3F, 60 cells; 4M, 74 cells); THC (4F, 59 cells; 4M, 87 cells); CBD (3F, 59 cells; 4M, 76 cells); THC+CBD (3F, 56 cells; 3M, 67 cells). Statistical significance was assessed using multilevel modeling. Data are presented as mean ± SEM derived from bootstrapping output; *p < 0.05; **p < 0.01; ***p < 0.001.

Total length, total branching points, and Sholl analysis, commonly used parameters [95], were analyzed to determine which pCB affects astrocyte branch patterns (Figure 3B-D). Our data show that the total length and the number of branching points are significantly increased in CBD-PCE male progenies, while increasing trends are observed in female progenies (Figure 3B-C). THC- and THC+CBD-PCE did not significantly impact astrocyte primary branch patterns in the mPFC-PrL L5 region. Sholl analysis revealed a significant increase in branch intersections following CBD-PCE in both males and females (Figure 3D).

### CBD-PCE results in significant sex-dependent changes in astrocyte fine processes

The significant increases in astrocyte major branch patterns induced by CBD-PCE prompted us to examine astrocyte fine processes, namely leaflets and endfeet [95]. To visualize these astrocytic processes, astrocytes were sparsely labeled with membrane-bound GFP by intraventricular injection of AAV5.GfaABC1D.PI.Lck-GFP.SV40 into CBD-PCE and vehicle control pups at P0, and brains were collected at P21 (Figure 4A) for 3D volume reconstructions and morphological analysis of individual GFP-labeled astrocytes in the mPFC-PrL L5/L6 (Figure 4B-D).

**Figure 4.**
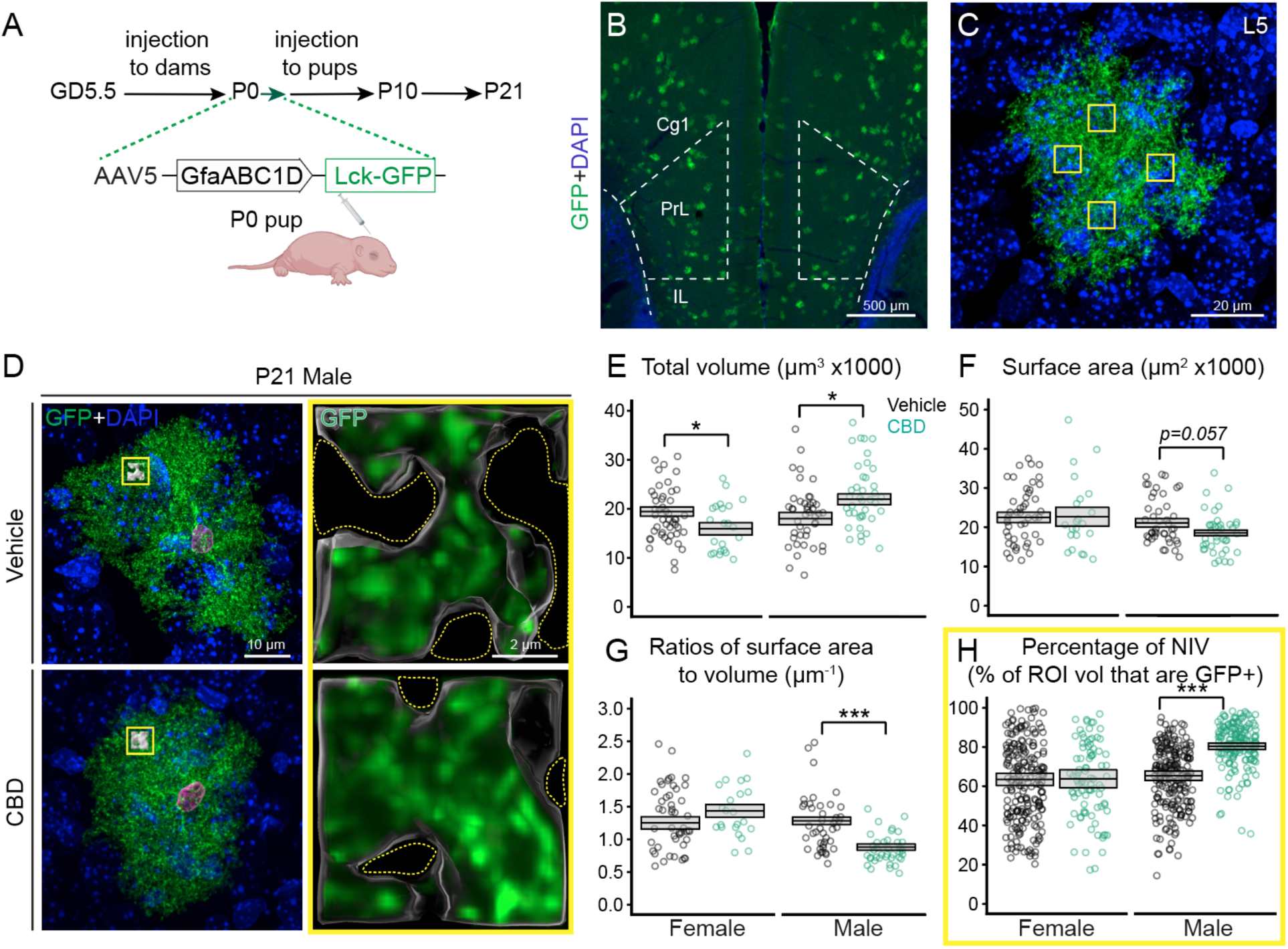
Detailed 3D morphological analysis of astrocytes labeled with membrane GFP. (A) Schematic showing P0 intraventricular injections with AAV5.GfaABC1D.PI.Lck-GFP.SV40 during the PCE treatment regimen. (B) Representative image of Lck-GFP-labeled astrocytes in mPFC-PrL (dashed lines). (C) High-magnification view of a representative Lck-GFP-labeled astrocyte located in L5. Yellow boxes indicate selected regions for NIV measurements. (D) Representative pairs of projected confocal image stacks of astrocytes from vehicle and CBD-PCE mPFC. Astrocyte nuclei were labeled with magenta. Yellow boxes indicate selected regions for NIV measurements. Yellow dashed lines denote regions within the ROI devoid of GFP signal. (E-H) Summary graph of total volume (E), surface area (F), the ratio of surface area-to-volume (G) and % of NIV in selected neuropil regions (H) (e.g. yellow boxes in C). For E-G, numbers of mice and cells analyzed per group: Vehicle (16F, 53 cells; 10M, 53 cells); CBD (6F, 23 cells; 10M, 43 cells). For H, numbers of mice and NIV analyzed per group: Vehicle (16F, 209 NIV; 10M, 211 NIV); CBD (6F, 89 NIV; 10M, 172 NIV). Statistical significance was assessed using multilevel modeling. Data are presented as mean ± SEM derived from bootstrapping output; *p < 0.05; **p < 0.01; ***p < 0.001. (E-G) Each data point represents one astrocyte. For NIV analysis summarized in (H), each data point corresponds to an ROI.

Whole-cell astrocyte volume analysis revealed that CBD-PCE resulted in significant sex-opposite changes in total cell volume: a decrease in females, while an increase in males (Figure 4E). No differences were detected in CBD-PCE astrocyte surface area, except for a trending decrease in males (Figure 4F). Astrocyte swelling has been observed in many pathological conditions and can be sensitively detected by a decrease in the surface-to-volume ratio [42–46]. Notably, our analysis shows that the surface area-to-volume ratios of CBD-PCE male astrocytes, but not female astrocytes, are significantly decreased compared to vehicle control astrocytes (Figure 4G).

Stogsdill et al. [56] have shown that astrocyte morphogenesis is linked to synaptogenesis by demonstrating changes in astrocyte fine membrane processes in the neuropil, quantified as neuropil infiltration volume (NIV), that correlate with neural circuit changes. To examine whether CBD-PCE alters NIV, we selected GFP-positive regions away from the astrocyte soma and main branches to quantify NIV in these identified neuropil regions (Figure 4C; see Supplementary Figure 4A-B for NIV segmentation workflow). Excitingly, we found that astrocyte NIVs are significantly higher in male CBD-PCE progenies than in control progenies, whereas no change was observed in females (Figure 4H).

### CBD-PCE increases AQP4 and Kir4.1 within astrocytic fine processes

The reduction in astrocyte surface area-to-total volume ratios in male CBD-PCE progenies suggests an alteration in the regulation of water and ion movements, as observed in previous studies [61, 96, 97]. Astrocyte fine process swelling could also explain the significant increases in NIV observed after CBD-PCE. AQP4, a water channel, and Kir4.1, an inwardly rectifying potassium channel, are enriched in astrocyte processes where they regulate water and ion flux across astrocytic membranes and help maintain extracellular ion homeostasis [45, 96, 98–101].

To examine the abundance of AQP4 and Kir4.1 and their association with astrocytic fine processes, we conducted AQP4 and Kir4.1 immunostaining with brain sections prepared from GfaABC1D.PI.Lck-GFP AAV-labeled brains after CBD-PCE or vehicle treatment (Figure 5). We found a significant increase in AQP4 abundance in the mPFC-PrL L5/L6 region of CBD-PCE male brains, but not female brains (Figure 5A-B). To further examine AQP4 in astrocytic processes, we used the Imaris Spots module to quantify AQP4 localized on astrocytic-GFP-positive processes. This analysis showed that astrocytic AQP4 abundance is significantly higher in male CBD-PCE mPFC compared to controls (Figure 5C). Similar analyses were conducted with Kir4.1. Here, the abundance of astrocytic Kir4.1 is also significantly increased in CBD-PCE male mPFC compared to control controls, while total Kir4.1 density remains unchanged (Figure 5D-F).

**Figure 5.**
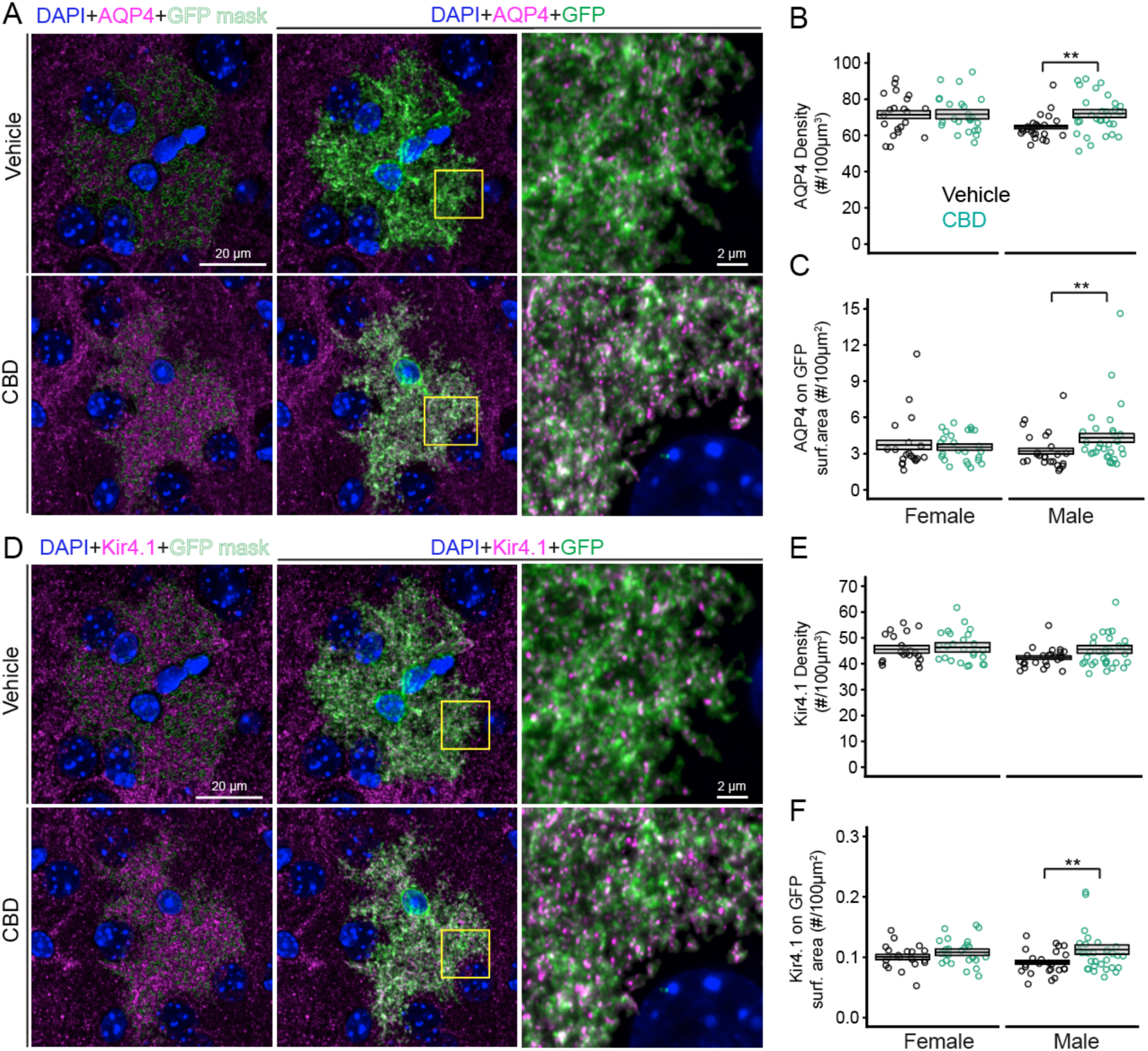
Quantification of astrocytic membrane AQP4 and Kir4.1. (A) Representative images acquired from males showing AQP4 (magenta) and astrocyte-GFP membrane rendered by Imaris on the left panel, while the corresponding confocal images are placed in the middle panel. ROIs highlighted in yellow boxes are shown at high magnification in the right panel. (B-C) Summaries for total AQP4 density (B) and the density of AQP4 located on GFP^+^ astrocyte membranes (C). (D) Representative images acquired from males showing Kir4.1 (magenta) and astrocyte-GFP membrane rendered by Imaris (left) from the same astrocytes shown in A. (E-F) Summaries for total Kir4.1 density (E) and the density of Kir4.1 on GFP^+^ astrocyte membranes (F). Numbers of experimental mice and images per group: Vehicle (6F, 21 images; 6M, 24 images); CBD (6F, 25 images; 10M, 31 images). Statistical significance was assessed using multilevel modeling. Data are presented as mean ± SEM derived from bootstrapping output; *p < 0.05; **p < 0.01; ***p < 0.001.

### CBD-PCE exerts differential impacts on intra- and thalamocortical tripartite glutamatergic synapses in a sex-dependent manner

Astrocyte fine processes in the neuropil area often participate in forming tripartite synapses [102–104]; conversely, synaptic activity itself can influence astrocyte morphology [97, 105, 106]. The observed increases in NIVs after CBD-PCE motivated us to assess whether CBD-PCE altered the density of tripartite synapses. Using high-resolution confocal imaging, we immuno-stained brain sections prepared from GfaABC1D.PI.Lck-GFP AAV-labeled brains after CBD-PCE or vehicle treatment with the antibody combination of VGLUT1+PSD95+GFP [107], VGLUT2+PSD95+GFP [62, 63], or VGAT+Gephrin+GFP [64, 65], to quantitatively examine the abundance of intracortical excitatory, thalamocortical excitatory, and inhibitory synapses, respectively.

We focused the analyses on tripartite synapses located in NIVs within the mPFC-PrL L5/6 region. Our data revealed a pronounced sex difference in the densities of cortical excitatory synapses in NIVs, with females containing significantly more synapses than males (Figure 6A-B). With respect to treatment effects, CBD-PCE significantly reduced intracortical excitatory tripartite synapse densities in females but not males. The analysis of thalamocortical tripartite synapses revealed that CBD-PCE significantly increases density in both females and males (Figure 6C-D), while having no impact on the inhibitory synapse densities in NIV (Figure 6E-F).

**Figure 6.**
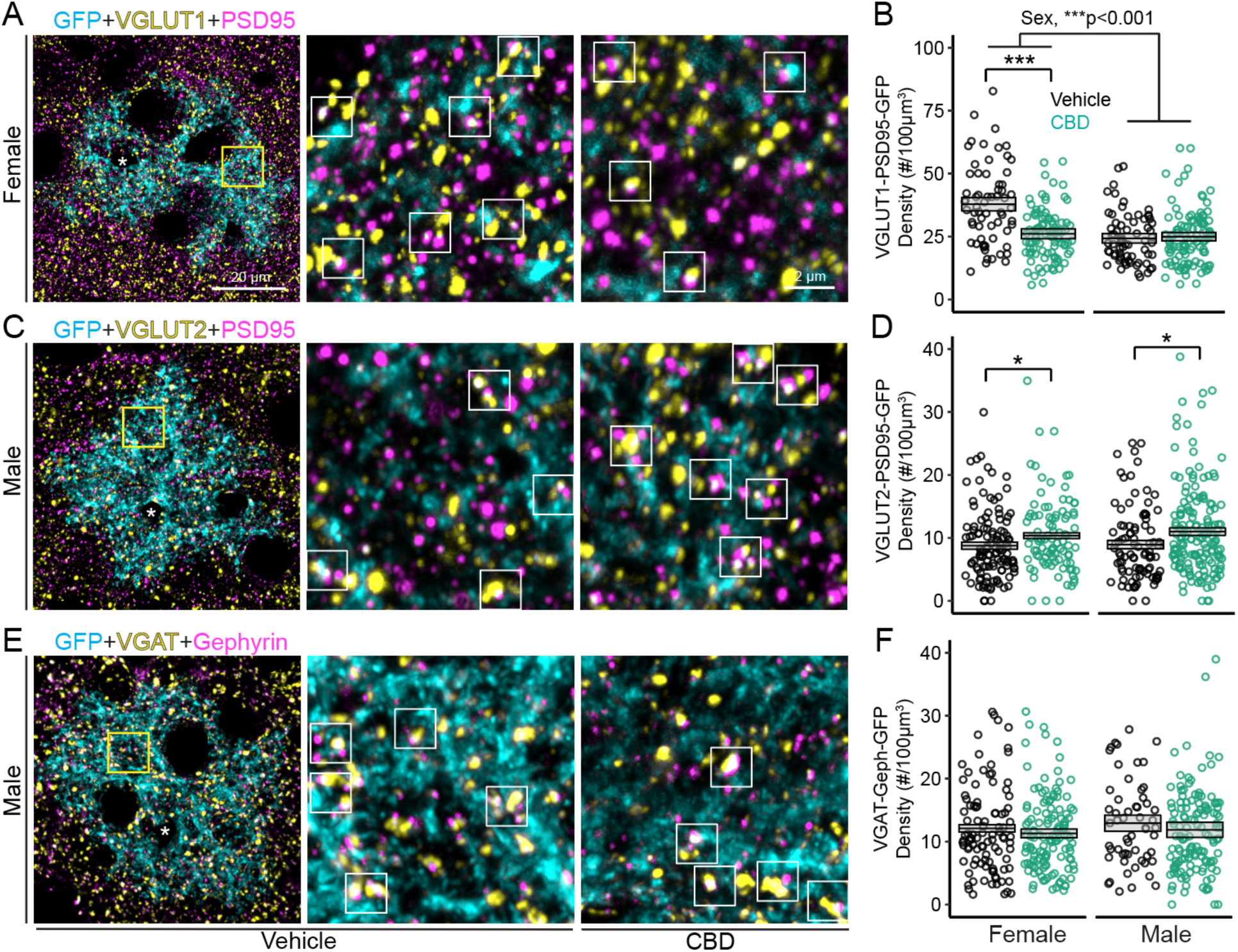
High-resolution image analysis of intracortical and thalamocortical and GABAergic tripartite synapses in mPFC-PrL. (A) Representative images showing the distribution of GFP-labeled astrocyte membranes (cyan), VGLUT1 (yellow) and PSD95 (magenta) in mPFC-PrL from P21 vehicle or CBD female brains. ROI highlighted with a yellow box is shown at high-magnification in the middle panel. White square boxes highlight synaptic puncta positive for VGLUT1 and PSD95 that are located within 0.5 µm of the GFP-labeled astrocyte membrane. (B) Summary for female and male VGLUT1+PSD95+GFP densities. (C) Similar arrangement as in (A) but for GFP (cyan)+VGLUT2 (yellow)+PSD95 (magenta) signals in males. (D) Summary of VGLUT2+PSD95+GFP densities. (E) Representative GFP (cyan)+VGAT (yellow)+Gephyrin (magenta) images with same arrangement as in A and C. (F) Summary for VGAT+Gephrin+GFP densities. Numbers of experimental mice, images and NIV analyzed per group: VGLUT1+PSD95 - Vehicle (4F, 16 images, 64 NIV; 4M, 17 images, 68 NIV); CBD (6F, 24 images, 96 NIV; 7M, 29 images, 104 NIV). VGLUT2+PSD95 - Vehicle (5F, 30 images, 120 NIV; 4M, 23 images, 92 NIV); CBD (4F, 24 images, 96 NIV; 7M, 40 images, 160 NIV). VGAT+Gephyrin - Vehicle (6F, 25 images, 100 NIV; 3M, 14 images, 56 NIV); CBD (7F, 32 images, 128 NIV; 7M, 33 images, 132 NIV). Statistical significance was assessed using multilevel modeling. Data are presented as mean ± SEM derived from bootstrapping output; *p < 0.05; **p < 0.01; ***p < 0.001. Each data point represents one ROI positioned within NIV (3-4 NIV per image).

Regarding total synaptic puncta density in a similar imaging volume used for tripartite synapse evaluations, we found significantly higher intracortical excitatory synapse densities in females than males for both the control and CBD-PCE groups (Supplementary Fig. 2A-B). Our analysis on thalamocortical synapses revealed a significant increase in CBD-PCE males but not females (Supplementary Fig. 2C-D). No difference in inhibitory synapse densities was identified between CBD-PCE or male vs female (Supplementary Fig. 2E-F). Together, our data suggest that CBD-PCE impacts astrocytic and synaptic compositions.

## DISCUSSION

In this study, we developed a preclinical PCE model to better mimic human cannabis exposure by accounting for mice being born with a less mature brain than humans [108, 109]. In addition, we optimized the dosing regimen to provide consistent pCB exposure throughout fetal brain development across all three trimesters of pregnancy. We then employed this pCB-PCE paradigm to evaluate effects on mPFC cytoarchitecture and found minimal changes in the distribution of neurons and glia. However, CBD- and THC+CBD-PCE significantly reduced astrocyte density in the mPFC, but only in male offspring. The most prominent reduction was found in the L5/L6 region of the mPFC, known for its critical roles in mediating top-down control and executive function [110–112]. Quantitative evaluation of astrocyte branch patterns in this brain region revealed significant increases in total length and branch number in male CBD-PCE progenies. Excitingly, our detailed morphological analysis with sparsely membrane-bound GFP-labeled astrocytes reveals that CBD-PCE significantly increases astrocyte total volumes and NIVs. This was accompanied by increases in AQP4 and Kir4.1 abundance, which likely mediate the reductions in surface-to-volume ratios, consistent with astrocytic process swelling.

Additionally, we found that CBD-PCE increases the density of tripartite synapses containing thalamocortical inputs in both sexes, while reducing intracortical excitatory tripartite synapse density only in females. In contrast, GABAergic tripartite synapse density remained unchanged in mPFC L5/L6 in both males and females. Taken together, we find that CBD-PCE exerts significant effects on astrocyte numbers and morphology predominantly in male progeny, while changing the composition of tripartite glutamatergic thalamocortical synapses in both sexes. This pCB-PCE model is designed to expose the developing embryonic and postnatal mouse brain with consistent pCB concentrations comparable to circulating plasma levels reported in moderate human cannabis users [113–116]. Here we show that the s.c. THC+CBD injections to the nursing dam result in pup brain levels of CBD that are about 20% of E18.5 levels (Supplementary Fig. 1). This suggests poor absorption of CBD from the pup gut, as CBD is present in mouse breast milk [9]. Thus, our model differs significantly from previous translational CBD models by injecting pregnant dams either by the exposure duration [17, 18] or the consistency of CBD concentration in the brains of the progeny [8]. Astrocyte proliferation and morphogenesis in mice occur mostly after birth [30, 117]. Thus, using previously established CBD-PCE paradigms, it is unlikely that CBD, via maternal exposure, will be present to exert impacts on astrocytes in the progeny’s brain. Conducting additional studies with this new CBD-PCE paradigm will help to determine if third-trimester CBD exposure produces lasting behavioral changes in adult progenies.

This study focused on determining the effects of pCB-PCE on astrocyte morphology, given their critical roles in brain function [29, 118]. We found CBD- but not THC-PCE alters astrocyte morphogenesis, with changes most pronounced in males. The increase in branching points and total astrocyte volume in the mPFC of male CBD-PCE progenies could be caused by one or more of the following mechanisms: (1) a direct influence of CBD on astrocyte patterning through CBD receptor(s); (2) neural circuit changes caused by exogenous CBD and adaptive alterations of astrocyte morphology; (3) increase in the number of branches to compensate for fewer astrocytes to maintain individual astrocyte tiling. The last scenario aligns with previous observations that astrocytes can dynamically adjust territory size and process elaboration in response to changes in local cellular composition or circuit demand [119, 120].

AQP4, the predominant water channel in the brain, is expressed in astrocyte endfeet and plays an important role in brain edema following ischemia [121]. Astrocytes swell in response to elevations in extracellular K^+^, as potassium uptake by astrocytes generates an osmotic gradient that drives water influx [122]. Accordingly, AQP4 is critical for astrocyte water movement and volume regulation [45, 46, 96, 98], and astrocyte swelling has been observed in response to changes in ionic environment or neuronal activity levels [44, 123, 124]. Kir4.1 is an inwardly rectifying K^+^ channel predominantly expressed in CNS glial cells [43], and often enriched alongside AQP4 in specific astrocytic membrane domains [42, 58]. Kir4.1 has been implicated in extracellular K^+^ homeostasis, maintenance of astrocyte resting membrane potential, cell volume regulation, and facilitation of glutamate uptake [43]. In this study, we found significant increases in AQP4 and Kir4.1 levels on astrocytic membranes labeled by Lck-GFP in the mPFC L5/6 of males after CBD-PCE. Taken together with increased astrocyte volume, these findings suggest that CBD-PCE alters astrocytic mechanisms involved in water and ion homeostasis, including processes related to K^+^ handling. Such changes could have important consequences for glial physiology and downstream effects on neurotransmission. Future studies will be important to examine how synaptic transmission is altered following CBD-PCE.

In addition to altering astrocyte morphology, we find that CBD-PCE also affects synaptic density and composition. In the mPFC-PrL L5/6, we found that there were more thalamocortical tripartite synapses in CBD-PCE male progenies, without changes in intracortical or inhibitory tripartite synapses. Interestingly, there are significantly fewer intracortical tripartite synapses in the mPFC-PrL L5/L6 of CBD-PCE female progenies. Overall, our data reveal morphological alterations in tripartite synapse composition by CBD-PCE, with input- and sex-specific effects. The observed differences may have resulted from differential sex developmental trajectories for the mPFC limbic region [125, 126]. Collectively, our findings underscore the importance of considering sex-specific astrocyte developmental trajectories in models of PCE and provide a foundation for understanding how early exogenous CBD exposure can impact astrocyte-synapse structural interface.

## Supporting information

Antibody table

## ACKNOWLEDGEMENTS

This work was supported by the following grants: National Institutes of Health grants DA053746 (HCL, KM) and P30DA056410 (HCL, HB, KM). We thank Scott Barton, Dr. Patricia Franco, Dr. Zhen Xian Niou, and Dr. Laszlo Barna for their technical support and Drs. István Katona and Norbert Hájos for their insightful discussion. We also thank Dr. Ben Deneen for providing us Aldh1l1-GFP mice.

## DATA AVAILABILITY STATEMENT

All data supporting the findings of this study are available from the corresponding author upon reasonable request. Processed dataset and analysis scripts can be shared for academic and noncommercial purposes.

---

**Supplementary Figure 1.**
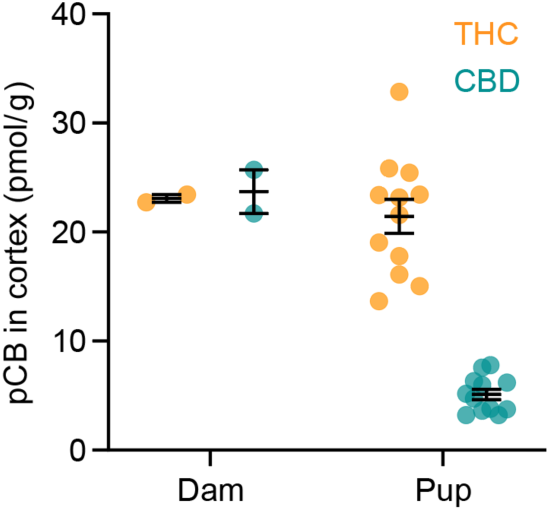
Comparative pCB levels in cortex of dams and pups at P10. Summary graph showing pCB levels in the cortices from P10 pup and their dams following 3 mg/kg THC+CBD s.c. to dams from six days after delivery for four days. Samples collected 2 hrs after last injection for MS measurements. Data are presented as mean ± SEM. P10 pup numbers: n=12; dams: n=2.

**Supplementary Figure 2.**
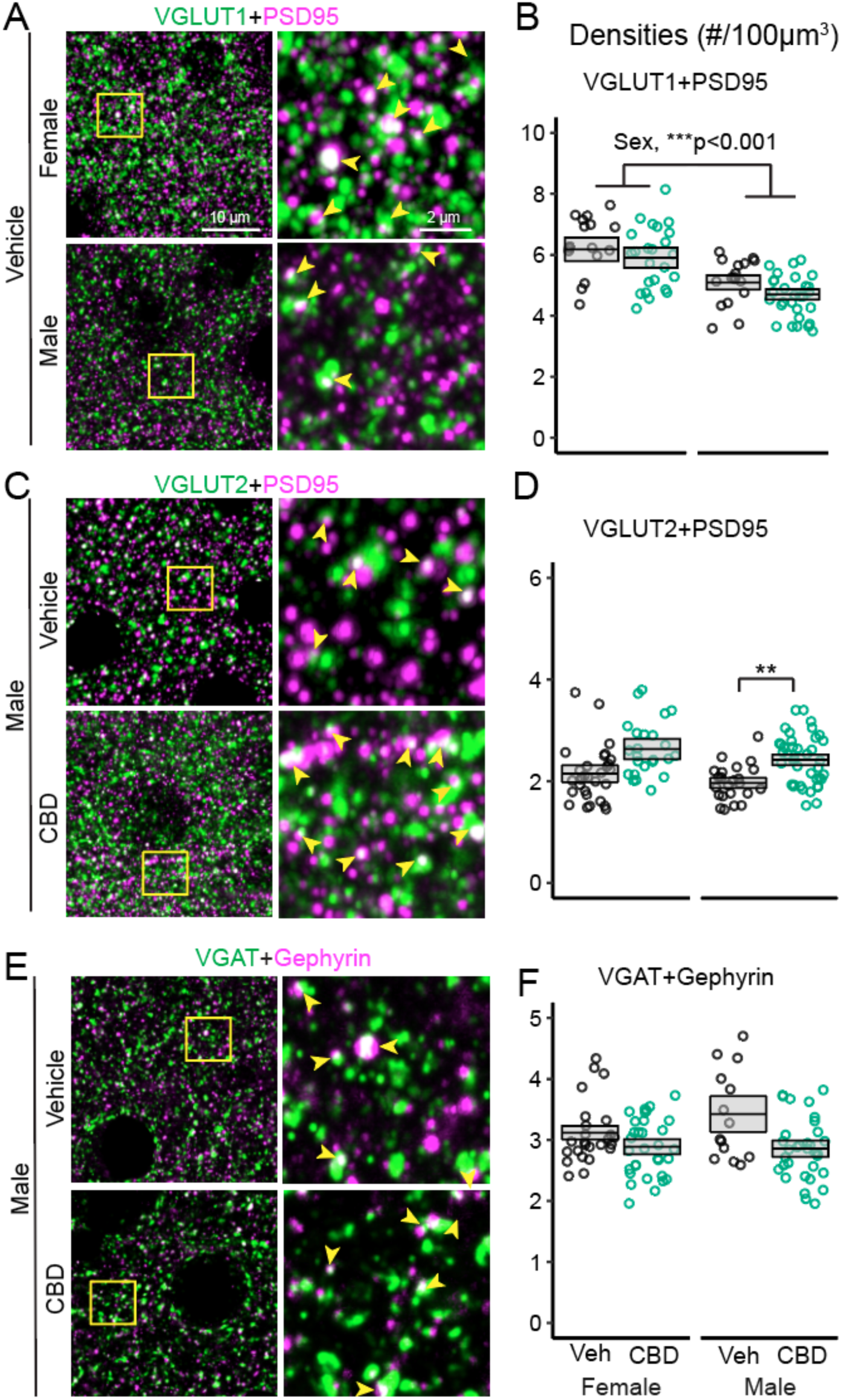
Synaptic puncta analysis for intracortical, thalamocortical and GABAergic synapses in vehicle and CBD-PCE mPFC-PrL at P21. (A) Representative projected images from confocal stacks for VGLUT1 (green) and PSD95 (magenta) staining from vehicle-treated male and female. High-magnification views of the yellow box indicated regions are shown on the right panels. Yellow arrowheads indicate VGLUT1+PSD95 juxtaposed synaptic boutons (white signals). (B) Summary for the density of VGLUT1+PSD95 juxtaposed synaptic sites. (C) Representative projected images from confocal stacks for VGLUT2 (green)+PSD95 (magenta) staining from vehicle and CBD males. High-magnification views of the yellow-boxed regions are shown in the right panels. Yellow arrowheads indicate VGLUT2+PSD95 juxtaposed synaptic boutons (white signals). (D) Summary of density of VGLUT2+PSD95 juxtaposed synaptic sites. (E) Representative projected images from confocal stacks for VGAT (green) and Gephyrin (magenta) staining from vehicle and CBD males. High-magnification views of the yellow box indicated regions are shown on the right panels. Yellow arrowheads indicate VGAT+Gephyrin juxtaposed synaptic boutons (white signals). (F) Summary of the densities of VGAT+Gephyrin juxtaposed synaptic sites. Numbers of experimental mice and images analyzed per group: VGLUT1+PSD95 - Vehicle (4F, 16 images; 4M, 17 images); CBD (6F, 24 images; 7M, 29 images). VGLUT2+PSD95 - Vehicle (5F, 30 images; 4M, 23 images); CBD (4F, 24 images; 7M, 40 images). VGAT+Gephyrin - Vehicle (6F, 25 images; 3M, 14 images); CBD (7F, 32 images; 7M, 33 images). Statistical significance was assessed using multilevel modeling. Data are presented as mean ± SEM derived from bootstrapping output; *p<0.05; **p<0.01; ***p<0.001.

**Supplementary Figure 3.**
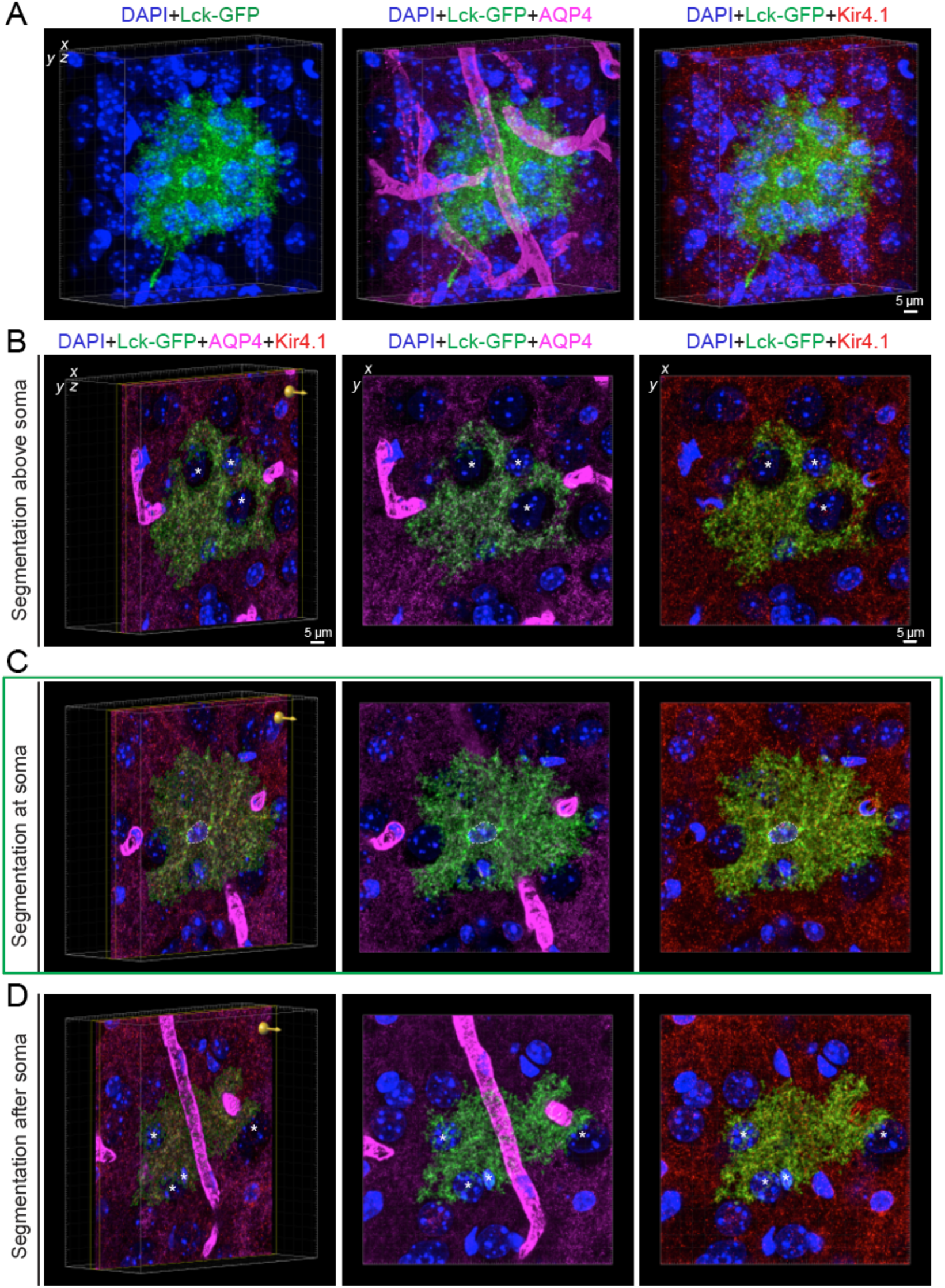
Representative panel images illustrate high-resolution whole astrocyte morphology exploration to determine AQP4 or Kir4.1 distributions. (A) Representative 3D view from Imaris encompassing a complete astrocyte co-stained with DAPI (left), AQP4 (middle), and Kir4.1 (right). The 42 µm z-stack was acquired at 0.1 µm z-step intervals with 3x software zoom and 2x frame averaging. The image was deconvolved prior to visualization. (B) Qualitative visualization reveals that this single astrocyte contacts at least two capillaries and exhibits punctate AQP4 and Kir4.1 expression along the astrocyte membrane across three segmented z-planes. Segmentation of 5 µm z-stack (yellow line boundaries) located above the astrocyte soma (B), showing consistent staining patterns for both markers. (*) Indicates nuclei of neighboring cells contained within the astrocyte territorial domain. (C) Segmentation of 5 µm z-stack including the astrocyte soma. Dotted lines area representing astrocyte nucleus. (D) Segmentation of 5 µm z-stack below the astrocyte soma where a major blood vessel is enwrapped in astrocyte membrane.

**Supplementary Figure 4.**
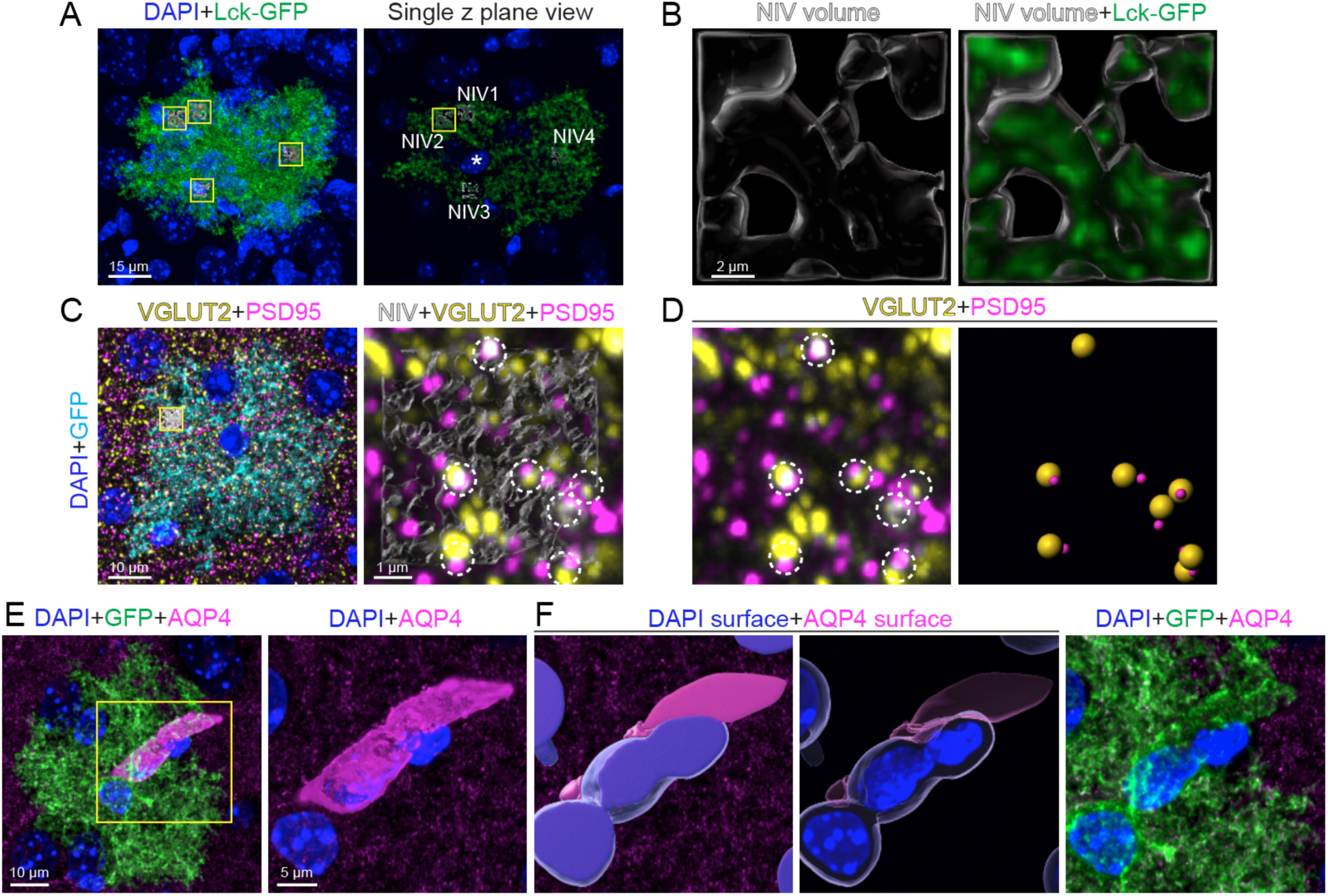
Graphic workflow for NIV analysis, object-based synaptic density within NIV and AQP4 vessel masking. (A) Representative image of whole-cell GFP-labeled astrocyte membrane and 4 distinct ROIs (yellow squares, left) representing each NIV selected within the z range (right). (B) Each NIV volume was rendered based on a fixed ROI size. Glass-like representation of GFP signal rendered with Imaris surface module (left) corresponding to GFP membrane signal occupying the inside of the ROI (right). Values were normalized as % of GFP signal occupying ROI total volume. (C) Representative four channel staining for DAPI, GFP-membrane (cyan) and pre- and postsynaptic markers (yellow and magenta, respectively). Yellow square ROI represents rendered NIV (white) (left). Representation of NIV as glass-like object rendered from membrane signal corresponding to the astrocyte membrane (right). Dotted line circles represent juxtaposed synaptic marker signals within 0.5 µm from each other and GFP surface, corresponding to potential tripartite site. (D) Representative images of pre- and postsynaptic markers signal within 0.5 µm of each other (left) and matching objects (right). (E) Representative image for astrocyte membrane (GFP), astrocyte endfeet AQP4 (magenta) perivascular structure (left). Zoomed in view of yellow ROI highlighting the structure to be processed in later steps (right). (F) Surface module rendering based on machine learning foreground and background training module on Imaris was applied to render DAPI and AQP4 surfaces (blue and magenta) (left). Transparent view of the signal contained within the surfaces (middle). These structures were masked out of the total image, removing the vasculature-like signal but preserving spots-cluster like AQP4 channel marker (right).

## REFERENCES

1. Spindle, T.R., M.O. Bonn-Miller, and R. Vandrey, Changing landscape of cannabis: novel products, formulations, and methods of administration. Curr Opin Psychol, 2019. 30: p. 98–102.

2. Wallis, D., et al., Predicting Self-Medication with Cannabis in Young Adults with Hazardous Cannabis Use. Int J Environ Res Public Health, 2022. 19(3).

3. Brown, Q.L., et al., Trends in Marijuana Use Among Pregnant and Nonpregnant Reproductive-Aged Women, 2002-2014. JAMA, 2017. 317(2): p. 207–209.

4. Volkow, N.D., et al., Self-reported Medical and Nonmedical Cannabis Use Among Pregnant Women in the United States. JAMA, 2019. 322(2): p. 167–169.

5. Young-Wolff, K.C., et al., Cannabis Use During Early Pregnancy Following Recreational Cannabis Legalization. JAMA Health Forum, 2024. 5(11): p. e243656.

6. Ochiai, W., et al., Maternal and Fetal Pharmacokinetic Analysis of Cannabidiol during Pregnancy in Mice. Drug Metab Dispos, 2021. 49(4): p. 337–343.

7. Moss, M.J., et al., Cannabis use and measurement of cannabinoids in plasma and breast milk of breastfeeding mothers. Pediatr Res, 2021. 90(4): p. 861–868.

8. Maciel, I.S., et al., Perinatal CBD or THC Exposure Results in Lasting Resistance to Fluoxetine in the Forced Swim Test: Reversal by Fatty Acid Amide Hydrolase Inhibition. Cannabis Cannabinoid Res, 2022. 7(3): p. 318–327.

9. Johnson, C.T., et al., Cannabinoids accumulate in mouse breast milk and differentially regulate lipid composition and lipid signaling molecules involved in infant development. BBA Adv, 2022. 2.

10. Harkany, T., et al., The emerging functions of endocannabinoid signaling during CNS development. Trends Pharmacol Sci, 2007. 28(2): p. 83–92.

11. Lu, H.C. and K. Mackie, Review of the Endocannabinoid System. Biol Psychiatry Cogn Neurosci Neuroimaging, 2021. 6(6): p. 607–615.

12. Bara, A., et al., Cannabis and synaptic reprogramming of the developing brain. Nat Rev Neurosci, 2021. 22(7): p. 423–438.

13. Abuhasira, R., L. Shbiro, and Y. Landschaft, Medical use of cannabis and cannabinoids containing products - Regulations in Europe and North America. Eur J Intern Med, 2018. 49: p. 2–6.

14. Dickson, B., et al., Recommendations From Cannabis Dispensaries About First-Trimester Cannabis Use. Obstet Gynecol, 2018. 131(6): p. 1031–1038.

15. Bhatia, D., et al., Cannabidiol-Only Product Use in Pregnancy in the United States and Canada: Findings From the International Cannabis Policy Study. Obstet Gynecol, 2024. 144(2): p. 156–159.

16. Stella, N., THC and CBD: Similarities and differences between siblings. Neuron, 2023. 111(3): p. 302–327.

17. Iezzi, D., et al., In utero exposure to cannabidiol disrupts select early-life behaviors in a sex-specific manner. Transl Psychiatry, 2022. 12(1): p. 501.

18. Iezzi, D., et al., Sex-specific disruptions in the developmental trajectory of anxiety-like behaviors due to prenatal cannabidiol exposure. Transl Psychiatry, 2025. 15(1): p. 354.

19. Laprairie, R.B., et al., Cannabidiol is a negative allosteric modulator of the cannabinoid CB1 receptor. Br J Pharmacol, 2015. 172(20): p. 4790–805.

20. Straiker, A., et al., Cannabidiol Inhibits Endocannabinoid Signaling in Autaptic Hippocampal Neurons. Mol Pharmacol, 2018. 94(1): p. 743–748.

21. Russo, E.B., et al., Agonistic properties of cannabidiol at 5-HT1a receptors. Neurochem Res, 2005. 30(8): p. 1037–43.

22. Martinez-Aguirre, C., et al., Cannabidiol Acts at 5-HT(1A) Receptors in the Human Brain: Relevance for Treating Temporal Lobe Epilepsy. Front Behav Neurosci, 2020. 14: p. 611278.

23. Alexander, C., et al., CBD and the 5-HT1A receptor: A medicinal and pharmacological review. Biochem Pharmacol, 2025. 233: p. 116742.

24. Bonnin, A., et al., Expression mapping of 5-HT1 serotonin receptor subtypes during fetal and early postnatal mouse forebrain development. Neuroscience, 2006. 141(2): p. 781–794.

25. Hanswijk, S.I., et al., Gestational Factors throughout Fetal Neurodevelopment: The Serotonin Link. Int J Mol Sci, 2020. 21(16).

26. Liu, J. and J.M. Lauder, Serotonin promotes region-specific glial influences on cultured serotonin and dopamine neurons. Glia, 1992. 5(4): p. 306–17.

27. Gaspar, P., O. Cases, and L. Maroteaux, The developmental role of serotonin: news from mouse molecular genetics. Nat Rev Neurosci, 2003. 4(12): p. 1002–12.

28. Aguado, T., et al., The endocannabinoid system promotes astroglial differentiation by acting on neural progenitor cells. J Neurosci, 2006. 26(5): p. 1551–61.

29. Khakh, B.S. and B. Deneen, The Emerging Nature of Astrocyte Diversity. Annu Rev Neurosci, 2019. 42: p. 187–207.

30. Akdemir, E.S., A.Y. Huang, and B. Deneen, Astrocytogenesis: where, when, and how. F1000Res, 2020. 9.

31. Abbink, M.R., et al., The involvement of astrocytes in early-life adversity induced programming of the brain. Glia, 2019. 67(9): p. 1637–1653.

32. Guayasamin, M., et al., Early-life stress induces persistent astrocyte dysfunction associated with fear generalisation. Elife, 2025. 13.

33. Tabata, H., Diverse subtypes of astrocytes and their development during corticogenesis. Front Neurosci, 2015. 9: p. 114.

34. Clavreul, S., et al., Cortical astrocytes develop in a plastic manner at both clonal and cellular levels. Nat Commun, 2019. 10(1): p. 4884.

35. Bushong, E.A., M.E. Martone, and M.H. Ellisman, Maturation of astrocyte morphology and the establishment of astrocyte domains during postnatal hippocampal development. Int J Dev Neurosci, 2004. 22(2): p. 73–86.

36. Farhy-Tselnicker, I. and N.J. Allen, Astrocytes, neurons, synapses: a tripartite view on cortical circuit development. Neural Dev, 2018. 13(1): p. 7.

37. Octeau, J.C., et al., An Optical Neuron-Astrocyte Proximity Assay at Synaptic Distance Scales. Neuron, 2018. 98(1): p. 49–66 e9.

38. Institoris, A., et al., Astrocytes amplify neurovascular coupling to sustained activation of neocortex in awake mice. Nat Commun, 2022. 13(1): p. 7872.

39. Chai, H., et al., Neural Circuit-Specialized Astrocytes: Transcriptomic, Proteomic, Morphological, and Functional Evidence. Neuron, 2017. 95(3): p. 531–549 e9.

40. Lanjakornsiripan, D., et al., Layer-specific morphological and molecular differences in neocortical astrocytes and their dependence on neuronal layers. Nat Commun, 2018. 9(1): p. 1623.

41. Thompson, A., et al., Brain-wide circuit-specific targeting of astrocytes. Cell Rep Methods, 2023. 3(12): p. 100653.

42. Nagelhus, E.A., T.M. Mathiisen, and O.P. Ottersen, Aquaporin-4 in the central nervous system: cellular and subcellular distribution and coexpression with KIR4.1. Neuroscience, 2004. 129(4): p. 905–13.

43. Nwaobi, S.E., et al., The role of glial-specific Kir4.1 in normal and pathological states of the CNS. Acta Neuropathol, 2016. 132(1): p. 1–21.

44. Reed, M.M. and B. Blazer-Yost, Channels and Transporters in Astrocyte Volume Regulation in Health and Disease. Cell Physiol Biochem, 2022. 56(S2): p. 12–30.

45. Pham, C., et al., Astrocyte aquaporin mediates a tonic water efflux maintaining brain homeostasis. Elife, 2024. 13.

46. Bakketun, C.B., et al., The Impact of Aquaporin-4 Deletion on K(+)-Induced Astrocytic Swelling Depends on K(+) Concentration. Glia, 2026. 74(1): p. e70086.

47. Aldinger, K.A., et al., Genetic variation and population substructure in outbred CD-1 mice: implications for genome-wide association studies. PLoS One, 2009. 4(3): p. e4729.

48. Cahoy, J.D., et al., A transcriptome database for astrocytes, neurons, and oligodendrocytes: a new resource for understanding brain development and function. J Neurosci, 2008. 28(1): p. 264–78.

49. Huang, A.Y., et al., Region-Specific Transcriptional Control of Astrocyte Function Oversees Local Circuit Activities. Neuron, 2020. 106(6): p. 992–1008.e9.

50. Torrens, A., et al., Comparative Pharmacokinetics of Delta(9)-Tetrahydrocannabinol in Adolescent and Adult Male Mice. J Pharmacol Exp Ther, 2020. 374(1): p. 151–160.

51. Leishman, E., et al., Cannabidiol’s Upregulation of N-acyl Ethanolamines in the Central Nervous System Requires N-acyl Phosphatidyl Ethanolamine-Specific Phospholipase D. Cannabis Cannabinoid Res, 2018. 3(1): p. 228–241.

52. Leishman, E., et al., Δ(9)-Tetrahydrocannabinol changes the brain lipidome and transcriptome differentially in the adolescent and the adult. Biochim Biophys Acta Mol Cell Biol Lipids, 2018. 1863(5): p. 479–492.

53. Paxinos, G., Keith B. J., The Mouse Brain in Stereotaxic Coordinates. 2008, San Diego, CA: Academic Press.

54. Pachitariu, M. and C. Stringer, Cellpose 2.0: how to train your own model. Nat Methods, 2022. 19(12): p. 1634–1641.

55. Kim, J.Y., et al., Intracerebroventricular viral injection of the neonatal mouse brain for persistent and widespread neuronal transduction. J Vis Exp, 2014(91): p. 51863.

56. Stogsdill, J.A., et al., Astrocytic neuroligins control astrocyte morphogenesis and synaptogenesis. Nature, 2017. 551(7679): p. 192–197.

57. Baldwin, K.T., et al., HepaCAM controls astrocyte self-organization and coupling. Neuron, 2021. 109(15): p. 2427–2442 e10.

58. Nagelhus, E.A., et al., Immunogold evidence suggests that coupling of K+ siphoning and water transport in rat retinal Muller cells is mediated by a coenrichment of Kir4.1 and AQP4 in specific membrane domains. Glia, 1999. 26(1): p. 47–54.

59. Verkman, A.S., et al., Aquaporin-4: orthogonal array assembly, CNS functions, and role in neuromyelitis optica. Acta Pharmacol Sin, 2011. 32(6): p. 702–10.

60. Smith, A.J. and A.S. Verkman, Superresolution Imaging of Aquaporin-4 Cluster Size in Antibody-Stained Paraffin Brain Sections. Biophys J, 2015. 109(12): p. 2511–2522.

61. Ciappelloni, S., et al., Aquaporin-4 Surface Trafficking Regulates Astrocytic Process Motility and Synaptic Activity in Health and Autoimmune Disease. Cell Rep, 2019. 27(13): p. 3860–3872 e4.

62. Fremeau, R.T., Jr., et al., The expression of vesicular glutamate transporters defines two classes of excitatory synapse. Neuron, 2001. 31(2): p. 247–60.

63. Hur, E.E. and L. Zaborszky, Vglut2 afferents to the medial prefrontal and primary somatosensory cortices: a combined retrograde tracing in situ hybridization study [corrected]. J Comp Neurol, 2005. 483(3): p. 351–73.

64. Chaudhry, F.A., et al., The vesicular GABA transporter, VGAT, localizes to synaptic vesicles in sets of glycinergic as well as GABAergic neurons. J Neurosci, 1998. 18(23): p. 9733–50.

65. Pizzarelli, R., et al., Tuning GABAergic Inhibition: Gephyrin Molecular Organization and Functions. Neuroscience, 2020. 439: p. 125–136.

66. Fogarty, M.J., et al., A method for the three-dimensional reconstruction of Neurobiotin-filled neurons and the location of their synaptic inputs. Front Neural Circuits, 2013. 7: p. 153.

67. Kuljis, D.A., et al., Fluorescence-Based Quantitative Synapse Analysis for Cell Type-Specific Connectomics. eNeuro, 2019. 6(5).

68. Simhal, A.K., et al., Multifaceted Changes in Synaptic Composition and Astrocytic Involvement in a Mouse Model of Fragile X Syndrome. Sci Rep, 2019. 9(1): p. 13855.

69. Zhou, B., et al., Astroglial dysfunctions drive aberrant synaptogenesis and social behavioral deficits in mice with neonatal exposure to lengthy general anesthesia. PLoS Biol, 2019. 17(8): p. e3000086.

70. Grecco, G.G., et al., Sex-Dependent Synaptic Remodeling of the Somatosensory Cortex in Mice With Prenatal Methadone Exposure. Adv Drug Alcohol Res, 2022. 2.

71. Haggerty, D.L., et al., Prenatal methadone exposure selectively alters protein expression in primary motor cortex: Implications for synaptic function. Front Pharmacol, 2023. 14: p. 1124108.

72. Kim, E. and M. Sheng, The postsynaptic density. Curr Biol, 2009. 19(17): p. R723–4.

73. Dumitriu, D., et al., Vamping: stereology-based automated quantification of fluorescent puncta size and density. J Neurosci Methods, 2012. 209(1): p. 97–105.

74. Irala, D., et al., Astrocyte-secreted neurocan controls inhibitory synapse formation and function. Neuron, 2024. 112(10): p. 1657–1675 e10.

75. Galbraith, S., J.A. Daniel, and B. Vissel, A study of clustered data and approaches to its analysis. J Neurosci, 2010. 30(32): p. 10601–8.

76. Aarts, E., et al., A solution to dependency: using multilevel analysis to accommodate nested data. Nat Neurosci, 2014. 17(4): p. 491–6.

77. de Melo, M.B., et al., Beyond ANOVA and MANOVA for repeated measures: Advantages of generalized estimated equations and generalized linear mixed models and its use in neuroscience research. Eur J Neurosci, 2022. 56(12): p. 6089–6098.

78. Huang, J.Y., et al., Protocol for detecting neuronal subnetworks from in vivo calcium imaging data using multiple clustering algorithms in MATLAB. STAR Protoc, 2025. 6(3): p. 104030.

79. Huang, J.Y., et al., From initial formation to developmental refinement: GABAergic inputs shape neuronal subnetworks in the primary somatosensory cortex. iScience, 2025. 28(3): p. 112104.

80. Workman, A.D., et al., Modeling transformations of neurodevelopmental sequences across mammalian species. J Neurosci, 2013. 33(17): p. 7368–83.

81. Zeiss, C.J., Comparative Milestones in Rodent and Human Postnatal Central Nervous System Development. Toxicol Pathol, 2021. 49(8): p. 1368–1373.

82. Gong, T., et al., A time-resolved multi-omic atlas of the developing mouse liver. Genome Res, 2020. 30(2): p. 263–275.

83. Knudsen, E.I., Sensitive periods in the development of the brain and behavior. J Cogn Neurosci, 2004. 16(8): p. 1412–25.

84. Vivi, E. and B. Di Benedetto, Brain stars take the lead during critical periods of early postnatal brain development: relevance of astrocytes in health and mental disorders. Mol Psychiatry, 2024. 29(9): p. 2821–2833.

85. Smith, A.M., et al., Effects of prenatal marijuana on visuospatial working memory: an fMRI study in young adults. Neurotoxicol Teratol, 2006. 28(2): p. 286–95.

86. Tortoriello, G., et al., Miswiring the brain: Delta9-tetrahydrocannabinol disrupts cortical development by inducing an SCG10/stathmin-2 degradation pathway. EMBO J, 2014. 33(7): p. 668–85.

87. Bara, A., et al., Sex-dependent effects of in utero cannabinoid exposure on cortical function. eLife, 2018. 7: p. e36234.

88. Willford, J.A., et al., A longitudinal study of the impact of marijuana on adult memory function: Prenatal, adolescent, and young adult exposures. Neurotoxicol Teratol, 2021. 84: p. 106958.

89. Swenson, K.S., et al., Fetal cannabidiol (CBD) exposure alters thermal pain sensitivity, problem-solving, and prefrontal cortex excitability. Mol Psychiatry, 2023. 28(8): p. 3397–3413.

90. Chen, H.T. and K. Mackie, Adolescent Delta(9)-Tetrahydrocannabinol Exposure Selectively Impairs Working Memory but Not Several Other mPFC-Mediated Behaviors. Front Psychiatry, 2020. 11: p. 576214.

91. Kenwood, M.M., N.H. Kalin, and H. Barbas, The prefrontal cortex, pathological anxiety, and anxiety disorders. Neuropsychopharmacology, 2022. 47(1): p. 260–275.

92. Mullen, R.J., C.R. Buck, and A.M. Smith, NeuN, a neuronal specific nuclear protein in vertebrates. Development, 1992. 116(1): p. 201–11.

93. Deloulme, J.C., et al., Nuclear expression of S100B in oligodendrocyte progenitor cells correlates with differentiation toward the oligodendroglial lineage and modulates oligodendrocytes maturation. Mol Cell Neurosci, 2004. 27(4): p. 453–65.

94. Du, J., et al., S100B is selectively expressed by gray matter protoplasmic astrocytes and myelinating oligodendrocytes in the developing CNS. Mol Brain, 2021. 14(1): p. 154.

95. Baldwin, K.T., K.K. Murai, and B.S. Khakh, Astrocyte morphology. Trends Cell Biol, 2024. 34(7): p. 547–565.

96. Benfenati, V., et al., An aquaporin-4/transient receptor potential vanilloid 4 (AQP4/TRPV4) complex is essential for cell-volume control in astrocytes. Proc Natl Acad Sci U S A, 2011. 108(6): p. 2563–8.

97. Toman, M., et al., The influence of astrocytic leaflet motility on ionic signalling and homeostasis at active synapses. Sci Rep, 2023. 13(1): p. 3050.

98. Haj-Yasein, N.N., et al., Glial-conditional deletion of aquaporin-4 (Aqp4) reduces blood-brain water uptake and confers barrier function on perivascular astrocyte endfeet. Proc Natl Acad Sci U S A, 2011. 108(43): p. 17815–20.

99. Nagelhus, E.A. and O.P. Ottersen, Physiological roles of aquaporin-4 in brain. Physiol Rev, 2013. 93(4): p. 1543–62.

100. Walch, E. and T.A. Fiacco, Honey, I shrunk the extracellular space: Measurements and mechanisms of astrocyte swelling. Glia, 2022. 70(11): p. 2013–2031.

101. Tyurikova, O., et al., Astrocyte Kir4.1 expression level territorially controls excitatory transmission in the brain. Cell Rep, 2025. 44(2): p. 115299.

102. Lawal, O., F.P. Ulloa Severino, and C. Eroglu, The role of astrocyte structural plasticity in regulating neural circuit function and behavior. Glia, 2022. 70(8): p. 1467–1483.

103. Salmon, C.K., et al., Organizing principles of astrocytic nanoarchitecture in the mouse cerebral cortex. Curr Biol, 2023. 33(5): p. 957–972 e5.

104. Benoit, L., et al., Astrocytes functionally integrate multiple synapses via specialized leaflet domains. Cell, 2025. 188(23): p. 6453–6472 e16.

105. Hirrlinger, J., S. Hulsmann, and F. Kirchhoff, Astroglial processes show spontaneous motility at active synaptic terminals in situ. Eur J Neurosci, 2004. 20(8): p. 2235–9.

106. Heller, J.P. and D.A. Rusakov, Morphological plasticity of astroglia: Understanding synaptic microenvironment. Glia, 2015. 63(12): p. 2133–51.

107. Wojcik, S.M., et al., An essential role for vesicular glutamate transporter 1 (VGLUT1) in postnatal development and control of quantal size. Proc Natl Acad Sci U S A, 2004. 101(18): p. 7158–63.

108. Clancy, B., et al., Extrapolating brain development from experimental species to humans. Neurotoxicology, 2007. 28(5): p. 931–7.

109. Cottam, N.C., et al., From Circuits to Lifespan: Translating Mouse and Human Timelines with Neuroimaging-Based Tractography. J Neurosci, 2025. 45(12).

110. Harris, K.D. and G.M. Shepherd, The neocortical circuit: themes and variations. Nat Neurosci, 2015. 18(2): p. 170–81.

111. Baker, A., et al., Specialized Subpopulations of Deep-Layer Pyramidal Neurons in the Neocortex: Bridging Cellular Properties to Functional Consequences. J Neurosci, 2018. 38(24): p. 5441–5455.

112. Anastasiades, P.G. and A.G. Carter, Circuit organization of the rodent medial prefrontal cortex. Trends Neurosci, 2021. 44(7): p. 550–563.

113. Blackard, C. and K. Tennes, Human placental transfer of cannabinoids. N Engl J Med, 1984. 311(12): p. 797.

114. Karschner, E.L., et al., Extended plasma cannabinoid excretion in chronic frequent cannabis smokers during sustained abstinence and correlation with psychomotor performance. Drug Test Anal, 2016. 8(7): p. 682–9.

115. Lucas, C.J., P. Galettis, and J. Schneider, The pharmacokinetics and the pharmacodynamics of cannabinoids. Br J Clin Pharmacol, 2018. 84(11): p. 2477–2482.

116. Millar, S.A., et al., A Systematic Review on the Pharmacokinetics of Cannabidiol in Humans. Front Pharmacol, 2018. 9: p. 1365.

117. Bayraktar, O.A., et al., Astrocyte development and heterogeneity. Cold Spring Harb Perspect Biol, 2014. 7(1): p. a020362.

118. Khakh, B.S., On astrocyte-neuron interactions: Broad insights from the striatum. Neuron, 2025. 113(19): p. 3079–3107.

119. Wilhelmsson, U., et al., Redefining the concept of reactive astrocytes as cells that remain within their unique domains upon reaction to injury. Proc Natl Acad Sci U S A, 2006. 103(46): p. 17513–8.

120. Khakh, B.S. and M.V. Sofroniew, Diversity of astrocyte functions and phenotypes in neural circuits. Nat Neurosci, 2015. 18(7): p. 942–52.

121. Papadopoulos, M.C. and A.S. Verkman, Aquaporin-4 and brain edema. Pediatr Nephrol, 2007. 22(6): p. 778–84.

122. Murakami, S. and Y. Kurachi, Mechanisms of astrocytic K(+) clearance and swelling under high extracellular K(+) concentrations. J Physiol Sci, 2016. 66(2): p. 127–42.

123. Mola, M.G., et al., The speed of swelling kinetics modulates cell volume regulation and calcium signaling in astrocytes: A different point of view on the role of aquaporins. Glia, 2016. 64(1): p. 139–54.

124. Lafrenaye, A.D. and J.M. Simard, Bursting at the Seams: Molecular Mechanisms Mediating Astrocyte Swelling. Int J Mol Sci, 2019. 20(2).

125. Premachandran, H., M. Zhao, and M. Arruda-Carvalho, Sex Differences in the Development of the Rodent Corticolimbic System. Front Neurosci, 2020. 14: p. 583477.

126. Knouse, M.C., et al., Sex differences in the medial prefrontal cortical glutamate system. Biol Sex Differ, 2022. 13(1): p. 66.

