## Supplementary material for "Perinatal cannabidiol exposure reshapes astrocyte morphology and tripartite synapse organization in a sex-dependent manner": Antibody table

**SUPPLEMENTAL TABLE:** List of antibodies used in this study

| Antibodies | Host/Reactivity | Source | Catalog # | Lot # | Dilutions |
|---|---|---|---|---|---|
| GFP (IgY) | Chicken | Aveslab | GFP-1020 | GFP917979 | 1:4000 |
| S100β | Rabbit | Abcam | ab52642 | 1068686-10 | 1:1000 |
| NeuN (IgG2a) | Mouse | Millipore | MAB377 | 2884594 | 1:1000 |
| VGLUT1 | Rabbit | Synaptic Systems | 135-303 | 4-109 | 1:2000 |
| VGLUT2 | Guinea pig | Synaptic Systems | 135-404 | 3-48 | 1:2000 |
| PSD95 (IgG1) | Mouse | Synaptic Systems | 124-011 | 1-44 | 1:1000 |
| VGAT | Rabbit | Synaptic Systems | 131-011 | 131011/51 | 1:2000 |
| Gephyrin | Mouse | Synaptic Systems | 147-021 | 147021/47 | 1:1000 |
| Aquaporin-4 | Mouse | Synaptic Systems | 429-011 | 1-4 | 1:1000 |
| Kir4.1 | Guinea pig | Synaptic Systems | 472-005 | 1-2 | 1:1000 |
| Alexa 488 | Goat x Chicken | Life Technologies | A11039 | 2566343 | 1:1000 |
| Alexa 555 | Goat x Mouse (IgG2a) | Life Technologies | A21137 | 2901506 | 1:1000 |
| Alexa 555 | Goat x Mouse (IgG1) | Life Technologies | A21127 | 2892443 | 1:1000 |
| Alexa 647 | Goat x Guinea pig | Life Technologies | A21450 | 2446026 | 1:1000 |
| Alexa 647 | Goat x Rabbit | Life Technologies | A21245 | 1752070 | 1:1000 |
